# Stonewall prevents expression of testis-enriched genes and binds to insulator elements in *D. melanogaster*

**DOI:** 10.1101/2021.03.18.435951

**Authors:** Daniel Zinshteyn, Daniel A Barbash

## Abstract

Germline stem cells (GSCs) are the progenitor cells of the germline for the lifetime of an animal. In *Drosophila*, these cells reside in a cellular niche that is required for both their maintenance (self-renewal) and differentiation (asymmetric division resulting in a daughter cell that differs from the GSC). The stem cell-daughter cell transition is tightly regulated by a number of processes, including an array of proteins required for genome stability. The germline stem-cell maintenance factor Stonewall (Stwl) associates with heterochromatin, but its molecular function is poorly understood. We performed RNA-Seq on *stwl* mutant ovaries and found significant derepression of many transposon families but not heterochromatic genes. We also discovered that testis-enriched genes, including the differentiation factor *bgcn* and a large testis-specific cluster on chromosome 2, are upregulated or ectopically expressed in *stwl* mutant ovaries. Surprisingly, we also found that RNAi knockdown of *stwl* in somatic S2 cells results in ectopic expression of these genes.

Using parallel ChIP-Seq and RNA-Seq experiments in S2 cells, we discovered that Stwl binds upstream of transcription start sites and localizes to heterochromatic sequences. We also find that Stwl is enriched at repetitive sequences associated with telomeres. Finally, we identify Stwl binding motifs that are shared with known insulator binding proteins. We propose that Stwl affects gene regulation by binding insulators and establishing chromatin boundaries.

## Introduction

Adult stem cells exist in tissues where there is constant turnover of cells, such as gonads where gametes are continually produced and released. Germline Stem Cells (GSCs) are one of several adult stem cell populations that inhabit the fruit fly ovary, residing in a niche environment that is required to maintain them (Song and Xie, 2002; Xie and Spradling, 2000). Stem cells undergo asymmetric cell division, resulting in one differentiated daughter cell and one daughter cell that is identical to the parent, thus undergoing self-renewal. For GSCs, the differentiated daughter cell is the cystoblast, which then undergoes four rounds of incomplete mitosis to form a 16-cell cyst. Upon completing these four rounds, one of the cystocytes in the 16-cell cyst enters meiosis while the other 15 undergo endoreduplication. The meiotic cell will differentiate into an oocyte while the other 15 will become nurse cells that provide maternal factors to the oocyte.

The entire germ cell population of the ovary is derived from the 2-3 GSCs in each germarium. *Drosophila* have an intricate regulatory network of factors that are required for normal GSC function (Xie, Ting, 2012), which can be broadly categorized as maintenance factors required for self-renewal or differentiation factors required for cystoblast production. Many of the genes involved in GSC regulation are pleiotropic for other functions inside and outside of the ovary. For example, Piwi is required for both GSC maintenance and differentiation (Lin and Spradling, 1997; Ma et al., 2014), as well as for silencing of transposable elements via the piRNA pathway (Klenov et al., 2011). Well-known differentiation factors include the translational repressors Bam, Bgcn, and Sxl. These proteins form a complex that represses mRNAs associated with GSC renewal, including the maintenance gene *nanos* (Li et al., 2013). Ovaries lacking in any of these differentiation factors exhibit a tumorous ovary phenotype, manifested as an overabundance of GSC-like cells that fail to become cystoblasts. Sxl is essential for the cell-autonomous sex determination of germ cells (Hashiyama et al., 2011)

The *stwl* gene was discovered in a *P*-element mutagenesis screen for female sterility and subsequently identified as a germline-expressed gene in an enhancer trap screen (Berg and Spradling, 1991; Karpen and Spradling, 1992). It is primarily expressed in germline cells of the ovary, with weak expression in GSCs and increased expression in GSC progenitor cells (cystoblasts) and beyond (Clark and McKearin, 1996). *stwl* mutant egg chambers contain 16 polyploid nurse cells, indicating that the cystocyte-to-oocyte transformation does not occur and that *stwl* is required for oocyte determination. Egg chamber growth in *stwl* null ovaries arrests between stages 4 and 7, with germ cells undergoing apoptosis. *stwl* is also required for GSC maintenance: mutant ovaries typically lack GSCs, especially in older flies (Akiyama, 2002). *stwl* mutant GSC clones are rapidly depleted from the ovary via differentiation into cystoblasts, and egg chambers derived from these clones exhibit oocyte determination defects as seen in *stwl* mutant animals (Maines et al., 2007).

Stwl is also involved in heterochromatin maintenance. *stwl* mutations are dominant suppressors of position-effect variegation, suggesting that Stwl is required to promote the spreading of heterochromatin (Maines et al., 2007). Stwl colocalizes with the heterochromatin-binding protein HP1a and dense, heterochromatin-like structures at the nucleolus in S2 cells, acts as a transcriptional repressor in *in vitro* experiments, and promotes the spreading of heterochromatic histone markers H3K9me3 and H3K27me3 in larvae (Yi et al., 2009).

Stwl also colocalizes with the insulator binding protein CP190 in terminal filament cells of the ovary and presents as puncti at the nuclear lamina (Rohrbaugh et al., 2013). Insulators are genomic regions that, when appropriately bound by insulator proteins, can prevent interaction between enhancers and their target promoters, or modulate the spreading of chromatin modifications (Dorman et al., 2007). Here, using a combination of transcriptional profiling in mutant ovaries and ChIP-Seq in S2 cells, we present evidence that Stwl has insulator binding properties.

## Methods

### Drosophila stocks

*P^{w[+mC]=lacW}^stwl^j6C3^* (*stwl^j6C3^*) was acquired from the Bloomington Stock Center [#12087]. This allele is female sterile and shows *stwl* mutant phenotypes (ovarian atrophy, loss of germline, lack of Orb accumulation in oocytes) when trans-heterozygous to a *stwl* deficiency chromosome (*Df(3L)Exel6122*) [Bloomington Stock Center #7601].

We found that the *stwl^j6C3^* chromosome is homozygous lethal, suggesting an accumulation of lethal recessive mutation(s). In order to remove these lethal mutations and homogenize the genetic background, we outcrossed *stwl^j6C3^* mutant females to males from an inbred *y w* strain (10 generations of inbreeding; strain will be subsequently referred to as *y w* F10) for 8 generations. Stocks were founded by balancing recombined 3rd chromosomes over a *TM6b* from *w*; *Sp*/*CyO*; *TM2*/*TM6b* stock in single female matings. Presence of the *P*-element insertion in *stwl^j6C3^* was followed by its *w^+^* marker and confirmed by PCR. The resultant stock produced viable and fertile homozygous males and viable but sterile homozygous females.

### Preparation of gonadal tissue for qRT-PCR and RNA-seq

All flies were raised at 25° C. Virgin males and females of each genotype were collected and aged for two days for the “older” samples; for the newly-eclosed samples, virgin females were collected and dissected immediately (<4 hours post-eclosion). Testis and ovary dissections were performed according to previously published protocols (Wong and Schedl, 2006; Zamore and Ma, 2011). Briefly, 15-30 flies at a time were sedated using CO_2_ and stored on ice. Gonads were extracted in ice-cold 1x PBS using sharp forceps, separated from gut tissue (and accessory glands, in males) and stored in ice-cold 1x PBS for ∼30 minutes. PBS was aspirated and tissues homogenized in 100-600 µl of Trizol (depending on total volume of dissected tissue) prior to snap-freezing in liquid NO_2_ and storage at −80° C. All sample replicates for RNA-seq consisted of ∼30 ovary/testis pairs, most of which were collected in single dissections at approximately the same time of day over a span of 23 days. Trizol homogenate from phenotypically “large” ovaries (2-day old *y w* F10) was diluted 1:10 prior to RNA extraction, to prevent overloading of columns.

RNA was extracted according to previously published protocols (Rio et al., 2010). Briefly, Trizol-homogenized tissue samples were thawed at room temperature and treated with 0.2 volumes chloroform to promote phase separation. RNA was extracted from the aqueous phase using Qiagen RNeasy Plus Mini Kit. This included application of aqueous phase to Qiagen gDNA Eliminator spin columns to limit carryover of genomic DNA. DNA contamination was also addressed by on-column DNAse digestion (Promega RQ1 DNAse). RNA quality and concentration was validated via Agilent Bioanalyzer; RNA quality for all samples was confirmed to have an RQN >=7.0 and at least 1.0 µg of starting material.

Stranded cDNA library preparation was performed by Polar Genomics (Ithaca, NY). mRNA was isolated and fragmented from total RNA pools, followed by 1st and 2nd (dUTP incorporated) strand synthesis. dsDNA was subsequently dA-tailed and adaptor-ligated, followed by size selection, UDG digestion to eliminate the second strand, and PCR amplification. All libraries (18 in total) were sequenced on a single lane of Illumina NextSeq (single-end, 75 bp).

During preliminary analyses of sequencing reads we found that 2 libraries (2 replicates from 0-day old *y w*; *stwl^j6C3^* ovaries) were of insufficient quality, likely due to contamination during sample recovery or library preparation. We therefore discarded these reads and prepared new samples. Ovaries were collected, dissected and homogenized in Trizol, as described above (with the exception that the ovary pool was increased from 30 to 45 ovary pairs per replicate). Stranded cDNA libraries were prepared as described above and subsequently sequenced on a single lane of Illumina HiSeq 2500 High Output (single-end, 50 bp).

For qRT-PCR, ovaries were dissected and processed as above, with three biological replicates of ∼30 ovary pairs each per sample. cDNA was synthesized with Invitrogen oligo-dT primers and reverse transcriptase using standard protocols. All qRT-PCR assays were performed with three technical replicates. Transcript abundance of each technical replicate was normalized to average levels of Rpl32 transcript in the source biological sample.

### Production and validation of polyclonal antibody against Stwl

A DNA fragment coding for amino acids 911-1037 from the C-terminus of the Stwl protein was amplified from *D. melanogaster* ovarian cDNA extracted from ∼20 *y w* F10 individuals. This region is lacking in predicted interaction domains, making it more likely to be accessible for immunoreactivity. The fragment was cloned into a N-terminal tagging MBP fusion vector (Genbank: AF097412.1) using NEB Gibson Assembly Kit (Sheffield et al., 1999) and transformed into chemically competent *E. coli* (One Shot^®^ TOP10). Successful assembly and transformation were confirmed via PCR and Sanger sequencing.

MBP-Stwl antigen was purified from induced bacterial culture using Amylose Resin (NEB: E8021L). Briefly, antigen expression was induced in 1 L of bacterial culture containing MBP-Stwl plasmid at log phase (OD 600 = 0.6) with 0.2 mM IPTG, then shaken for ∼18 hours at 18° C. Bacteria were pelleted and resuspended in lysis buffer (50 mM Tris pH 8.8, 200 mM NaCl, 2 mM DTT, 1 mM PMSF, 1 mg/ml lysozyme, 1x Roche cOmplete™ EDTA-free Protease Inhibitor Cocktail) at 4° C. Lysate was sonicated on ice to ensure thorough lysis, then spun at 20,000x g for 45 minutes to pellet debris. Supernatant was then applied to Amylose Resin on column. Stwl-MBP bound resin was washed 4 times with 1 column volume of low salt buffer (50 mM Tris pH 8.8, 200 mM NaCl, 2 mM DTT), followed by 4 washes with 1 column volume of high salt buffer (50 mM Tris pH 8.8, 1.5 M NaCl, 2 mM DTT) and another 4 washes with 1 column volume of low-salt buffer. Stwl-MBP was eluted with 10 mM maltose in low-salt buffer. Presence of 57.5 kDa MBP-stwl protein was confirmed using Coomassie stain on 10% SDS PAGE; concentration was estimated using Bradford assay. Protein-containing fractions were pooled using Amicon® Ultra-4 10K Centrifugal Filter Devices to a final concentration of 1.0 mg/ml.

Purified protein was submitted to Pocono Rabbit Farm & Laboratory Inc. for injection. Two guinea pigs (henceforth referred to as GP 76 and GP 77) were selected for antigen injection based on absence of background signal in pre-immune sera (determined by probing wild-type *D. melanogaster* ovaries with sera in immunofluorescence assays).

We found that both antibodies recognize a ∼135 kDa protein in wild-type ovaries (Supplementary Fig 1) and S2 cells (Fig 3A). This signal is absent in null and RNA knockdown samples, as well as wild-type lysates probed with pre-immune sera. The primary *stwl* transcript is predicted to produce a 112.9 kDa protein; previous work has shown that antibodies against Stwl recognize a similarly sized protein (Clark and McKearin, 1996).

We performed immunofluorescence (IF) experiments to confirm that the Stwl antibodies target a nuclear protein and to compare to IF experiments done with other Stwl antibodies. *D. melanogaster* ovaries were dissected from wild-type (*y w* F10) individuals in cold 1x PBS and fixed in 4% paraformaldehyde with 0.1% Triton X-100 in PBS. Tissue was then washed 3x in PBT (1x PBS with 0.1% Tween 20), followed by 4x washes in PBTA (PBT with 1.5% BSA). Samples were incubated overnight at 4°C with primary antibody at a concentration of 1:200 for Stwl antisera and 1:200 for Rabbit Vasa from Santa Cruz Biotechnology, Inc. Following 3x washes in PBT and 4x washes in PBTA, tissue was incubated for 2 hours with secondary antibodies (1:500 Goat anti-Guinea Pig Rhodamine Red-X, 1:500 Goat anti-Rabbit Alexa 488). Following 3x washes in PBT, tissue was mounted in vectashield with DAPI and imaged on a Zeiss Confocal. We found that both antibodies specifically labeled germ cell nuclei in testis and ovaries (Supplementary Fig 2). Furthermore, ectopic expression of HA-tagged Stwl [FlyORF stock #F001844] colocalized with signals from both Stwl antisera (Supplementary Fig 3) (Bischof et al., 2013).

### Cell Culture and RNAi

S2 cells were cultured in M3+BPYE medium, made as directed from Shields and Sang Powdered Medium (Sigma S-8398), supplemented with 0.5 g KHCO_3_, 1 g yeast extract and 2.5 g bactopeptone per liter, pH adjusted to 6.6 and sterile-filtered. 100x Antibiotic-Antimycotic (Thermo-Fisher 15240062) and Fetal Bovine Serum (Sigma F2442 Lot # 078K8405) were added to concentrations of 1x and 10%, respectively. Cells were maintained at 25°C and passaged every 3-4 days for 7 passages prior to use for RNAi experiments.

For dsRNA-induced knockdown, cells were plated in serum-free medium at a concentration of 2.5 million cells/ml, then treated with 30 µg/ml of *lacZ*- or *stwl*-dsRNA for 60 minutes before addition of M3/BPYE medium containing 13% FBS (final concentration, 10% FBS, 7.5 µg/µl dsRNA). Cells were chemically cross-linked and frozen after 3 days.

RNA was synthesized using NEB HiScribe™ T7 High Yield RNA Synthesis Kit (E2040S) from PCR products generated from YEp365 plasmid (*lacZ* control) or genomic DNA extracted from S2 cells. For efficient *stwl* KD we generated three distinct dsRNAs from reference, each targeting the second exon of *stwl*, which is present in all *stwl* transcripts (Hu et al., 2016). S2 cells were treated with 10 µg/ml of each dsRNA.

### Chromatin Immunoprecipitation in S2 cells

Subsequent to dsRNA treatment for 3 days, cells were centrifuged for 5 minutes at 1000xg followed by removal of media, washed once in 1x PBS, and resuspended in 1x PBS and cross-linked via addition of 16% paraformaldehyde to 1% final concentration for 2 minutes at room temperature. Cross-linking was quenched by addition of 2.5 M glycine in 1x PBS (final concentration 0.15 M) for 5 minutes at room temperature. Cells were nutated for 15 minutes at 4°C, spun and washed in 1x PBS brought to 4°C, and pelleted and flash-frozen in liquid nitrogen.

Cells were thawed on ice and lysed in RIPA buffer containing 0.1% SDS, 1% Nonidet P-40, and 1 tablet/10 ml Pierce™ Protease Inhibitor Mini Tablets, EDTA-free (A32955), for 20 minutes. Lysates were then sonicated at high intensity in a Bioruptor (Diagenode) water bath to shear DNA to desired size range (300-500 bp) for 45 minutes total with cycles of 20 seconds on, 1-minute rest, with quick spins of lysates every 15 minutes to settle samples and re-fill the Bioruptor with ice-cold water.

6 µl of freshly thawed Stwl antisera and pre-immune sera were added to 300 µl of cell lysate (1:50 dilution) and incubated overnight at 4°C. Cell lysates were prepared with approximately 34,000 cells per µl, so that each IP experiment was performed on roughly 10 x 10^6^ cells. IP complexes were immunoprecipitated with Invitrogen Dynabeads™ Protein A for Immunoprecipitation (10001D). Prior to use, beads were washed 2x 10 minutes in blocking buffer containing 1 mg/ml BSA, 1 mg/ml propyl vinylpyrrolidone blocking agent, and 1 tablet/10 ml Pierce™ Protease Inhibitor Mini Tablets, EDTA-free (A32955), and 1x 10 minutes in chilled RIPA buffer (also with protease inhibitor). IP samples were added to blocked beads and incubated at 4°C for 2 hours; 50 µl of beads were used for each IP.

Beads were washed 1x in low-salt buffer, 2x in high-salt buffer, 1x in LiCl buffer, and 2x in TE buffer. IP complexes were eluted from beads in 10% SDS, 1M NaHCO_3_ elution buffer for 30 minutes at 65°C. Cross-linking was reversed by addition of 5 M NaCl to 0.2 M NaCl final concentration and overnight incubation at 65°C. DNA was treated with RNAse A for 2 hours at 37°C and Proteinase K for 2 hours at 55°C, then cleaned using Qiagen QIAquick Gel Extraction Kit. Input samples were frozen following sonication, then thawed and reverse-crosslinked as above. DNA libraries were prepared using NEBNext Ultra II DNA Library Prep Kit for Illumina (NEB# E7645), using Ampure XP beads for cleanup, without size selection.

RNA was extracted in triplicate from S2 cells originating from the same populations used for ChIP-Seq experiments, using Qiagen RNEasy plus extraction kit, which includes additional elimination of gDNA from samples. All RNA samples had RQN>7.0, as determined by Bioanalyzer instrument (Agilent). cDNA libraries were prepared using NEBNext Ultra II Directional RNA Library Prep Kit for Illumina (E7760G), using Ampure XP beads for cleanup. All cDNA and ChIP libraries (22 in total) were pooled together and sequenced on a single lane of Illumina NextSeq (single-end, 75 bp). RNA-Seq data processing, QC, and analysis of S2 cell samples was performed as described in the “Read processing, alignment, and normalization” section.

### Read processing, alignment, and normalization

We assayed quality of raw reads in fastq format using FastQC (version 0.11.6) and trimmed reads for adapter sequences and quality using Trimmomatic (version 0.32); (java -jar trimmomatic-0.32.jar SE [raw_reads.fq] [trimmed_reads.fq] ILLUMINACLIP:TruSeq3-SE.fa:2:30:10 SLIDINGWINDOW:4:20 MINLEN:50 AVGQUAL:20) (Andrews, 2010; Bolger et al., 2014). We used FastQ Screen to identify non-*Drosophila* contaminants in our libraries (Wingett and Andrews, 2018).

For RNA-Seq, we aligned reads to a curated list of consensus sequences for repetitive elements using relaxed bowtie2 settings (bowtie2 -x [repetitive_consensus_sequences.fasta] -U [trimmed_reads.fq] -S [repetitive_alignment.sam] --un-gz [unmapped_reads.fq.gz] --score-min L,0,-1.5 -L 11 -N 1 -i S,1,.5 -D 100 -R 5). Unmapped reads from this alignment were saved and aligned to the unmasked *Drosophila melanogaster* genome (r6.03) using bowtie2 default settings (Langmead and Salzberg, 2012). We used a custom Perl script to count the number of reads aligning to repetitive sequences; we utilized HTSeq version 0.6.0 to count the number of reads aligning to exons in the genomic alignment (Anders et al., 2015). We concatenated the read counts into a single file for each sample.

In order to normalize for sequencing bias resulting from GC-content bias or batch effects, we normalized the read counts using EDAseq (Risso et al., 2011). We removed counts for all genes with a mean read count less than or equal to 10 across all samples. We then performed within-lane normalization for GC-content: read counts within individual samples were transformed via full-quantile normalization between feature strata to normalize for GC-content of assayed genes. Between-lane normalization was then performed (again using full-quantile normalization between feature strata) to account for differences in sequencing depth, and offset values were generated for each transcript in the count matrix so that raw counts could be analyzed for differential expression analysis.

In order to provide context for our biological observations, we compared *stwl* mutant ovary data to *bam*, *egg*, *wde*, *hp1a*, *Sxl*, and *piwi* mutant ovaries (Gan et al., 2010; Peng et al., 2016; Shapiro-Kulnane et al., 2015; Smolko et al., 2018). Alignment and normalization for all datasets were as described above.

For ChIP-Seq, trimmed reads were aligned to the unmasked *Drosophila melanogaster* genome (r6.03) using bowtie2 default settings (Langmead and Salzberg, 2012). All reads with mapping quality <20 were removed using SAMtools (Li et al., 2009b). All alignment files were corrected for GC bias using the deepTools commands computeGCbias and correctGCbias (Ramírez et al., 2014). Briefly, the distribution of GC content per read is assessed over the contents of each alignment file, typically revealing overamplification of high-GC content sequences. The correctGCbias command generates an alignment file identical to the original, except with reads artificially removed or duplicated at biased regions to eliminate GC bias.

### Repetitive DNA alignment and analysis for ChIP-Seq

Limitations and challenges of identifying enriched repetitive elements from ChIP-Seq data have been well documented (Huang et al., 2013; Lin et al., 2015; Marinov et al., 2015). With relatively short (75 bp) single-end reads, it is nearly impossible to identify the genomic origin of most reads coming from repetitive DNA, and therefore enrichment cannot be called against a true background signal. We therefore instead calculated differential enrichment of repetitive DNAs in IP samples relative to mock samples, normalized against genomic reads. The process is explained below in greater detail.

For analysis of repetitive DNA, reads were trimmed and aligned as described for RNA-Seq reads. For genomic reads, rather than counting reads aligned to gene bodies as we did for RNA-Seq analysis, we calculated the number of reads aligned to each 1 kb bin of the genome. We concatenated the repetitive and genomic read counts into a single file for each sample.

### Differential expression/enrichment analysis

We analyzed count data using DESeq2 (Love et al., 2014). We imported raw counts and offsets as described above from genes with mean read count >10 across all samples. For testis, we estimated differential expression of genes between mutant and wild-type samples. For ovary comparisons, age of the samples was taken into account: genes were called as differentially expressed if the normalized read counts from the experimental genotype were consistently different from those in the control genotype, excluding those cases where genes were found to be differentially expressed between 0- and 2-day old samples.

For ChIP-Seq, we estimated differential enrichment of genomic regions between IP and mock samples, taking into account the antibody used (GP 76 or GP 77) and the source S2 population (replicate 1 or 2). A genomic region was reported as differentially expressed only when the normalized read counts for that region were consistently greater in IP samples than in mock samples, and not due to differences in IP conditions (source animal antibody or source cell population). PCA on the count matrix confirmed that the majority of the variance in the count data was explained by the variance between mock and IP samples (Supplementary Figure 4).

Subsequent to differential expression/enrichment analysis, all log_2_(fold-change) estimates were transformed using apeGLM shrinkage estimator to reduce variability in LFC values among low-count genes (Zhu et al., 2018). Shrunken LFC values were used for all subsequent analyses, including overrepresentation tests, gene set enrichment analysis, and Gene Ontology analyses, implemented using the R package ClusterProfiler (Yu et al., 2012).

### ChIP-Seq peak calling and analysis

Each IP experiment was performed with 2 biological replicates; each biological replicate originated from a single 150 cm^2^ flask of *lacZ*-dsRNA-treated S2 cells. From each flask, we immunoprecipitated chromatin using Stwl antisera 76 and 77 (IP), pre-immune sera 76 and 77 (mock), and also sequenced input DNA. Therefore each ChIP-Seq experiment could be called against enrichment from its own input DNA, and mock datasets against both antibodies could be used to exclude spurious peaks.

We followed peak calling standards established by the ModERN consortium (Kudron et al., 2018). We performed peak calling on all mock and IP samples using the peak calling algorithm MACS2 (Feng et al., 2012). In order to generate an intentionally noisy set of peaks for downstream IDR (irreproducible discovery rate) analysis, peaks were called with low stringency (FDR<0.75) as follows: macs2 callpeak -t {IP/Mock.bam} -c {Input.Bam} -g 142573024 --tsize 75 -n output.file -m 2 50 -q 0.75 --keep-dup all. This command generates a large set of statistically insignificant peaks which can be fed into the IDR algorithm (Li et al., 2011). Confident peak sets were identified by performing IDR analysis on peaksets between biological replicates, using an IDR cutoff of 0.05; significant peaks passing this IDR threshold co-occur in the same genomic location at similar intensities. IDR was also done on mock samples with a much looser restriction (IDR<0.25), in order to create a more expansive list of peaks that could potentially be generated from biological noise (Supplementary Fig. 5).

After IDR, any peaks in the IDR 76 and IDR 77 IP peaksets that overlapped with spurious peaks from either mock were removed using BEDTools subtract (Quinlan and Hall, 2010). These filtered peaksets were then merged together using BEDTools merge so that the final Stwl peakset contained the union of peaks confidently called in IPs from either antibody. Finally, peak calling was repeated using MACS2 broadpeak setting (mfold 2-50, q<0.50), and the same steps were followed as before. The final broadpeak and narrowpeak calls were merged together to form a single set of peaks in broadpeak format. Motif identification, which requires narrow, sharply defined peaks, was done on only the narrowpeak calls; all other analyses were performed on the broadpeak format calls.

For motif analysis, we extracted summits from 1,379 narrowpeak calls as described above. Symmetrical peaks were then extracted as 500 bp sequences centered at each summit. These sequences were loaded onto the MEME-ChIP web browser (version 5.0.5) and motifs were identified using MEME, DREME, and CentriMo programs on default settings (Bailey et al., 2009).

### Identifying tissue-enriched and ectopically expressed genes

We utilized RPKM values from the modENCODE anatomy RNA-Seq dataset to classify all genes according to tissue-biased expression (Brown et al., 2014). The Tau metric is among the most simple and reliable tools for determining tissue-specific expression of a given gene (Kryuchkova-Mostacci and Robinson-Rechavi, 2017; Yanai et al., 2005). Tau was calculated from log_2_(RPKM) values in a subset of available tissues (Supplementary Tables 1,2). Tau values range from 0 to 1, corresponding to the range from ubiquitous expression to highly tissue-specific expression. Genes with Tau>= 0.7 were considered tissue-specific; to identify which tissue(s) each of these genes are enriched in, we assigned any tissue where log_2_(RPKM) is greater than 1.5 standard deviations above the mean log_2_(RPKM) across all tissues to that gene. According to this classification, we found that 45.5% of annotated genes exhibit tissue-specific expression, meaning that transcripts for those genes are preferentially enriched in one or more of the represented tissues (Supplementary Table 3).

Ectopic gene expression refers to the expression of a gene in a tissue where it is silent under normal conditions. Due to the nature of ectopic expression (i.e. an increase in transcript abundance from a baseline of very low counts), it is challenging to capture accurate log_2_(Fold-change) values, especially since GC-normalization typically requires removal of low-count genes. Ectopic gene expression in ovaries, testes, and S2 cells was assayed from library size-corrected RPKM values calculated in DESeq2, without the removal of low-count genes. We defined ectopic gene expression by 1) identifying the gene as phenotypically silent in wild-type tissue and 2) finding that gene expression increases significantly by at least 2-fold in the mutant tissue. Genes with mean RPKM < 2.0 in a given WT tissue according to a Mann-Whitney (p<0.1) were considered transcriptionally inert. Inert genes where mean_RPKM_null/mean_RPKM_WT>2.0 were subjected to a BH-corrected Mann-Whitney test (p<0.25) to identify ectopic expression.

## Results

### Stwl deficient ovaries exhibit TE derepression

A screen for genes whose RNAi-induced knockdown (KD) in ovaries leads to misexpression of TEs found that germline KD of *stwl* results in moderate derepression of three TE transcripts (*Het-A, blood, and burdock*), as determined by qRT-PCR (Czech et al., 2013). We performed our own qRT-PCR experiments to test for misexpression of the LTR retrotransposon *Copia*, and the non-LTR retrotransposons *Het-A, 412*, and *I* element. *Het-A, Copia* and *I* element are germline-restricted, while *412* is expressed in both germline and ovarian follicle cells (Li et al., 2009a; Malone et al., 2009). We tested for misexpression of these TEs in ovaries dissected from 2-day old trans-heterozygous null (*stwl^j6c3^/Df(3L)Exel6122*), hemizygous (*stwl^+^/Df(3L)Exel6122*), and wild-type (*stwl^+^/stwl ^+^*) flies. We found that each of these TEs is derepressed in trans-heterozygous null ovaries, relative to hemizygous and wild-type ovaries (Fig 1A).

**Figure 1.**
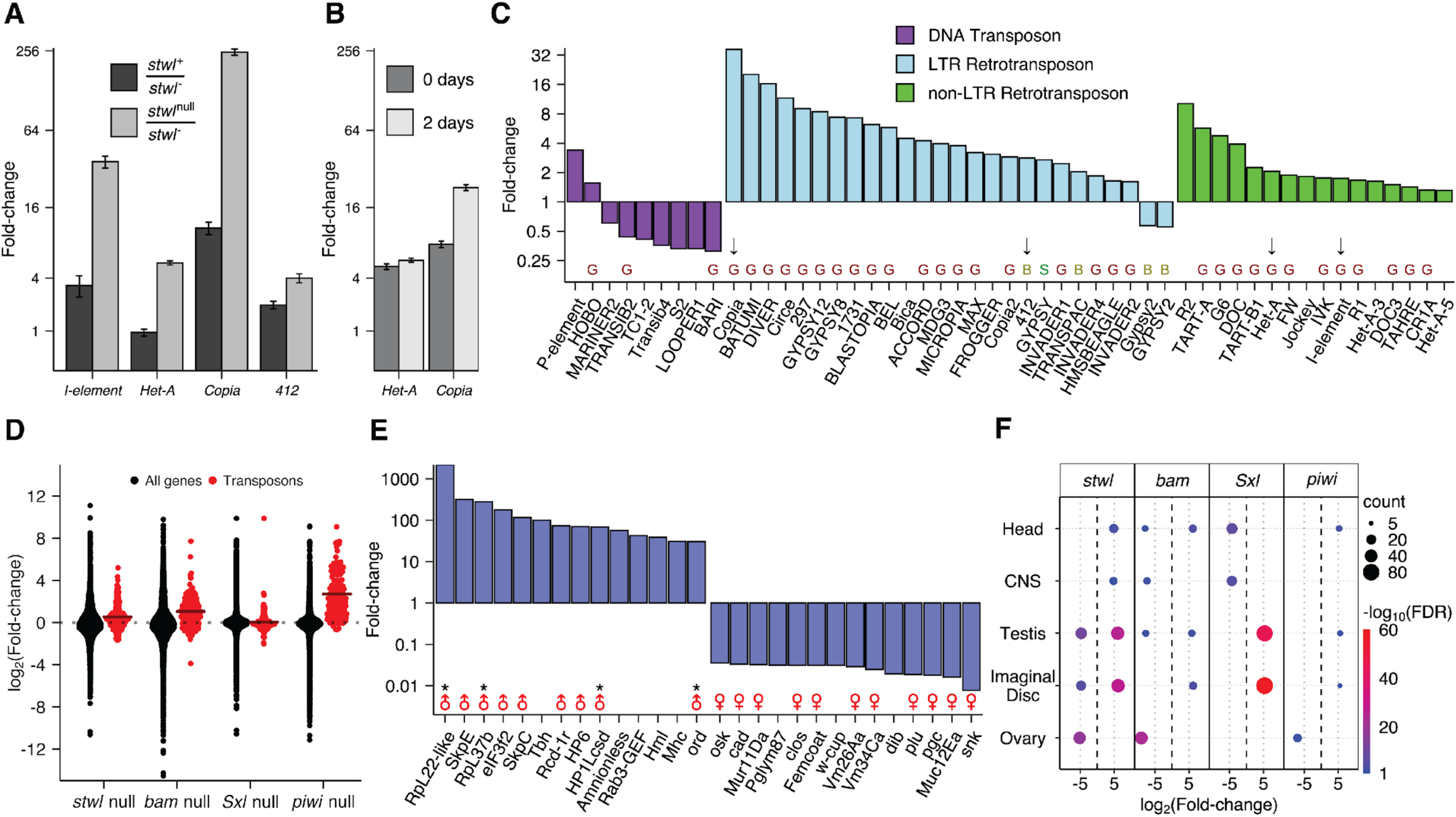
Loss of *stwl* results in upregulation of transposons and testis-enriched genes. (A) qRT-PCR of TEs from 2-day old ovaries, scaled to wild-type. *stwl*^null^ is a mutant allele (*stwl^j6c3^*), *stwl*^-^ is a deficiency allele (*Df(3L)Exel6122*). (B) TEs are upregulated in both 0- and 2-day old ovaries. qRT-PCR of TEs from *stwl* null (*stwl^j6c3^*/*stwl^j6c3^*) ovaries, scaled to *stwl*^+^/*Df(3L)Exel6122*. (A-B) Mean and SE plotted from 3 biological replicates, each with 3 technical replicates. (C) Fold-change of TEs in *stwl* null (*stwl^j6c3^*/*stwl^j6c3^*) relative to wild-type from RNA-Seq assay of 0- and 2-day old ovaries. Black arrows point to TEs validated with qRT-PCR data in Figs 1A and/or 1B. “G”,”S”,”B” indicates whether TE is typically expressed in germline, ovarian soma, or both, respectively (Malone et al., 2009). (D) log_2_Fold-change (LFC) of TEs vs. all genes from *stwl*, *bam*, *Sxl*, and *piwi* null ovaries, relative to wild-type. Crossbars show the mean LFC for all TEs. (E) Fold-change of the top 14 and bottom 14 most affected annotated genes (based on FlyBase annotations) in *stwl* null ovaries relative to wild-type. Male and female symbols mark genes with testis- and ovary-enriched wild-type expression, respectively; “*” marks genes that are part of the 59C4-59D testis-specific cluster described in Figure 2. (F) Enriched tissue classes among the top and bottom 1% of misregulated genes. Average LFC is plotted for each set of tissue-enriched genes enriched among *stwl* null ovaries relative to wild-type. Only gene sets with FDR <0.05 are plotted.

The *stwl* mutant phenotype presents challenges for interpreting assays of transcript abundance. *stwl* mutant ovaries are largely agametic as a consequence of GSC loss and defects in oocyte determination (Akiyama, 2002; Clark and McKearin, 1996; Maines et al., 2007). Nurse cells in *D. melanogaster* ovaries are polyploid and produce large quantities of mRNA that are maternally inherited by the developing oocyte. Differential expression between agametic mutant and wild-type ovaries might therefore reflect extensive differences in the cellular makeup of the ovaries rather than changes in transcript abundance specifically due to *stwl*. In order to account for differences in tissue composition, we chose the approach utilized by previous studies, which is to assay transcripts derived from extremely young ovaries (Shapiro-Kulnane et al., 2015; Sun and Cline, 2009). These authors reasoned that dissection of ovaries from newly-eclosed individuals (dissected within 24 hours of eclosion) would limit the amount of late-stage egg chambers and eggs that are present.

We assayed TE transcripts from newly-eclosed ovaries from both *stwl^j6c3^*/*Df(3L)Exel6122* and *stwl^+^*/*Df(3L)Exel6122*, as above. We found that the fold-increase of *Het-A* in trans-heterozygous nulls relative to hemizygotes is similar in newly-eclosed and 2-day-old ovaries (5-fold and 6-fold increase, respectively) (Fig 1B). *Copia* transcript is also derepressed in newly-eclosed *stwl* transheterozygous null ovaries (8-fold increase over hemizygotes), though this derepression phenotype is not as large as the one observed in 2-day-old ovaries (23-fold increase over hemizygotes). We conclude that the TE derepression phenotype we and Czech et al. (2013) observed is likely due to loss of *stwl* activity, not to a general loss of germ cells.

In order to identify the genome-wide consequences of *stwl* loss, we performed RNA-Seq on ovaries dissected from newly-eclosed and 2-day old wild-type (*y w*) and *stwl* null (*y w; stwl^j6c3^/stwl^j6c3^*) individuals. The goal of this experiment was to identify and classify genes and TEs which are consistently differentially expressed in *stwl* mutants. Therefore, we incorporated all four sample types (newly-eclosed wild-type, newly-eclosed mutant, two-day old wild-type, two-day old mutant) into a generalized linear model using DESeq2 (Love et al., 2014). A gene was only considered differentially expressed in *stwl* nulls if the transcript count for that gene significantly changed across both null samples relative to wild-type; that is, if the gene was differentially expressed between the two genotypes, regardless of age.

After accounting for potential batch effects and GC-content bias (see Methods), sample-to-sample distances for the resultant count matrices confirmed that the biological replicates for each sample type cluster together (Supplementary Fig 6). Principal Component Analysis (PCA) of the count data demonstrated that the samples are primarily stratified according to ovary maturity (Supplementary Fig 7). Principal Component 1 (PC1) accounts for 58% of the variance in the count matrix, which separates mature ovaries (2-day old *stwl^+^/stwl ^+^* wild-type) from immature ovaries (2-day old *stwl^j6c3^/stwl^j6c3^* null, 0-day *stwl^j6c3^/stwl^j6c3^* null, and 0-day old *stwl^+^/stwl ^+^*wild-type). These trends in the PCA support the rationale behind our experimental design, in that comparing null and WT ovaries at two time-points more accurately identifies genes that are differentially expressed due to genotype.

We found that 4,839 genes (out of 10,165 genes with mean count >10 across all ovary samples) are differentially expressed, with 2,147 genes upregulated in *stwl* null and 2,692 downregulated in *stwl* null (48%, 21%, and 26% of expressed genes) (Supplementary Fig 8). We also found that *P-element* transcript increases ∼4-fold in *stwl* null ovaries; this can be explained by the *P-element* insertion into the *stwl* locus that created the *stwl^j6c3^* allele and serves as an internal validation for the presence of the *stwl^j6c3^* allele. The RNA-Seq data showed that repetitive elements are strongly impacted by loss of *stwl* (Fig 1C). These repeats include the *Copia, Het-A, 412*, and *I element* elements we identified by qRT-PCR.

### Loss of Stwl, Bam, and Piwi, but not Sxl results in TE derepression

The terminal phenotype of *stwl* mutant ovaries is sterility caused by loss of germline stem cells and apoptosis of differentiated germ cells. DNA damage is also apparent in these sterile ovaries, possibly due to *stwl’*s requirement for maintenance of heterochromatin (Yi et al., 2009). It is also possible that TE derepression is a consequence of these defects, rather than reflecting a direct role of Stwl in TE silencing. To help distinguish between these possibilities, we analyzed published RNA-seq data generated from ovaries for mutations in various genes affecting GSC maintenance or differentiation. These included the differentiation factor *bag-of-marbles* (*bam*), the sex-determination master regulator *Sex-Lethal* (*Sxl*), and the GSC maintenance factor and piRNA targeting protein *piwi* (*piwi*). *bam* and *Sxl* deficient ovaries exhibit a “bag of marbles” phenotype that is characteristic of disruption of differentiation factors that results in over-proliferation of GSC-like cells (Gan et al., 2010; Shapiro-Kulnane et al., 2015; Smolko et al., 2018). *piwi* mutants exhibit GSC maintenance defects similar to those in *stwl* mutants (Cox et al., 1998; Klenov et al., 2011; Lin and Spradling, 1997; Peng et al., 2016). We found that loss of *bam* and *piwi*, but not *Sxl*, results in upregulation of TEs, particularly LTR retrotransposons and germline-expressed TEs (Fig 1D, Supplementary Fig 9). These results suggest that the *stwl* TE derepression phenotype is not an indirect reflection of the loss of GSCs, since it also occurs in *bam* mutants, which have the opposite phenotype of GSC overproliferation. It is also notable though that the magnitude of effects in *stwl* mutants is substantially lower than in *piwi* mutants, suggesting that *stwl* may not be a direct repressor of TEs as *piwi* is. We suggest instead that loss of *stwl* may lead to widespread but moderate derepression of TEs through its role in insulator function described below.

### A subset of testis-enriched genes are highly upregulated in *stwl* null ovaries

*Sxl* is required in ovaries for silencing of testis-specific transcripts (Shapiro-Kulnane et al., 2015). We tested whether *stwl*, *bam* and *piwi* mutant ovaries exhibit abnormal derepression of testis-specific genes. We utilized RPKM values from the modENCODE anatomy RNA-Seq dataset to classify all genes according to tissue-biased expression (Brown et al., 2014). We found that testis-enriched genes are among the most upregulated genes in *stwl* null ovaries, while ovary-enriched genes are among the most downregulated (Fig 1E-F).

While the apparent enrichment of testis-enriched transcripts at the top of the range of LFC values in *stwl* null ovaries might suggest that upregulation would also be detected by Gene Set Enrichment Analysis (GSEA), testis-enriched transcripts as a group are downregulated in *stwl* null ovaries (Supplementary Fig 10). This reflects a limitation in GSEA performance previously noted when attempting to perform analyses on large and potentially complex gene sets (Hong et al., 2014; Warden et al., 2013). The essential problem is that these sets include genes that are misregulated in both directions, presumably because members of the same pathway may be either down- or upregulated in response to misexpression of an upstream activator or suppressor. We therefore performed an over-representation test for tissue-enriched genes among the top and bottom 1% of expressed genes in each RNA-Seq experiment, to identify strong biases at the tips of the LFC ranges (Fig 1F). Our top/bottom percentile over-representation tests confirmed that testis-enriched genes are over-represented within the top 1% of most highly upregulated genes in agametic ovaries, but they are much more prevalent in *stwl* and *Sxl* null ovaries, relative to *bam* or *piwi* null ovaries.

We also examined the impact of *stwl* loss on the expression of genes normally enriched in non-gonadal tissues. Misexpression of imaginal-disc-enriched genes largely mirrors misexpression of testis-enriched genes, while genes enriched in adult head, and pharate and larval stage Central Nervous System (CNS) are also upregulated (Fig 1F).

### Loss of *stwl* results in ectopic expression of a testis-specific gene cluster

In order to test whether specific regions of the genome are misregulated, we plotted LFC by genomic location (Fig 2A). We found a striking pattern of expression at 59C4-59D on chromosome 2R, where 11 genes clustered within 227.5 Kb are strongly upregulated in *stwl* null ovaries. Four of the genes in this cluster are among the most strongly upregulated genes in *stwl* null ovaries (Fig 1E). Coexpressed gene clusters are common in many species, adding a dimension of organization to the genome by allowing groups of adjacent genes to be regulated simultaneously. Testis-specific gene clusters are particularly common in *D. melanogaster*, with 59C4-59D being the largest, and their expression is tightly regulated to prevent somatic expression (Shevelyov et al., 2009). We confirmed that the 59C4-59D cluster described by Shevelyov et al. is composed mostly (28/34 total genes) of testis-enriched genes, most of which are absent from the wild-type ovarian transcriptome (Supplementary Table 4).

**Figure 2.**
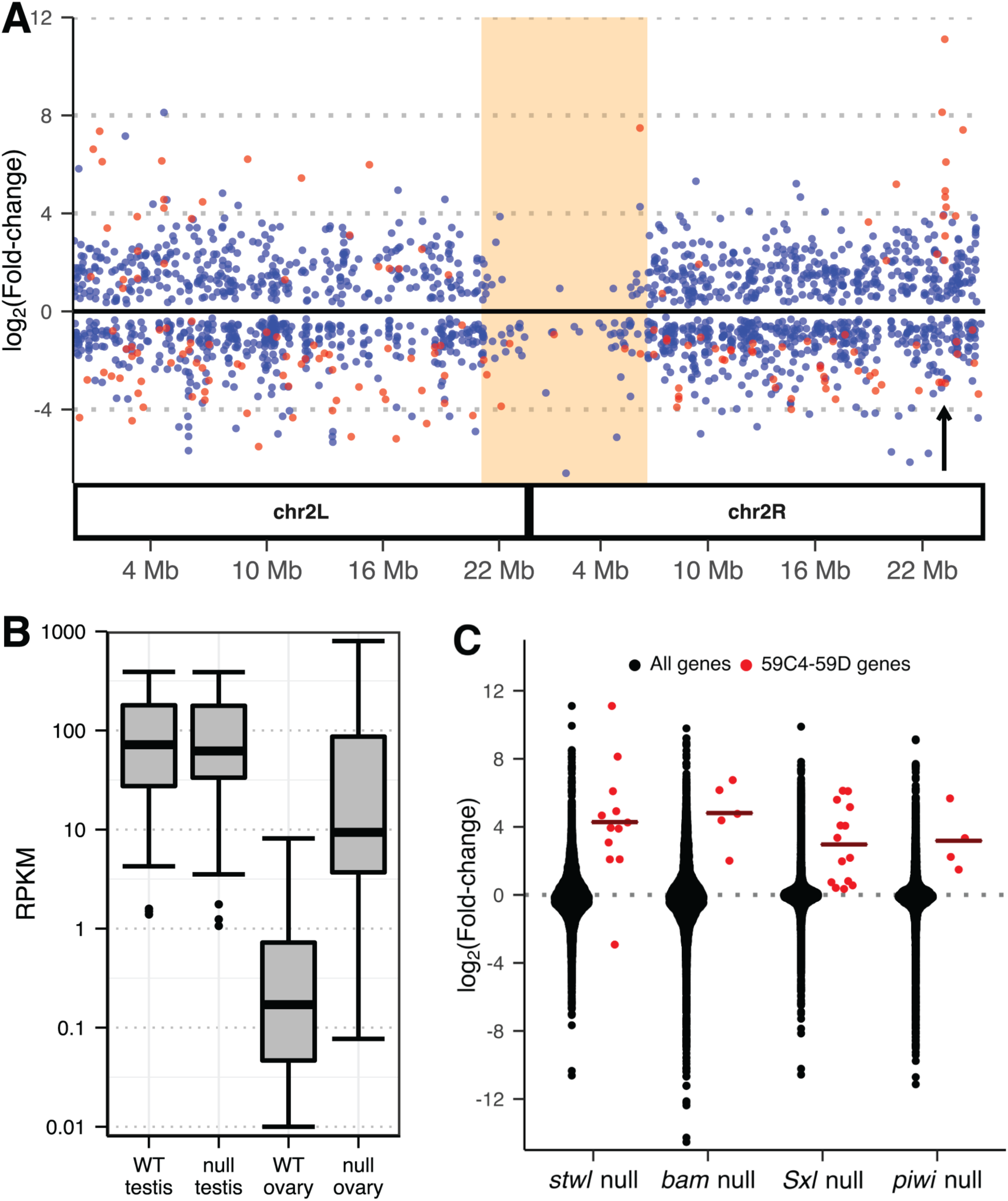
A cluster of testis-specific genes is derepressed in *stwl* null ovaries. (A) Differential expression (DE) of genes in ovaries (*stwl* null/WT) along chromosome 2. Only DE genes (FDR <0.01) are plotted. Testis-enriched genes are red (see Methods). Shaded orange area marks pericentromeric heterochromatin; arrow points to the testis-enriched cluster at 59C4-59D. (B) Reads Per Kilobase of transcript per Million mapped reads (RPKM) of genes in the 59C4-59D cluster in wild-type and *stwl* null gonads. Low-count outliers are not plotted. (C) log_2_Fold-change (LFC) of 59C4-59D cluster vs. all genes from *stwl*, *bam*, *Sxl*, and *piwi* null ovaries, relative to wild-type. Crossbars show the mean cluster LFC.

Loss of the H3K9me3 pathway components results in ectopic expression of testis-enriched genes (Shapiro-Kulnane et al., 2015; Smolko et al., 2018). Similarly, we found that the genes of the 59C4-59D cluster are transcriptionally inert in ovaries and become ectopically expressed in *stwl* null ovaries (Fig 2B, Supplementary Table 4). We also found that this cluster is upregulated in *bam*, *Sxl*, and *piwi* null ovaries (Fig 2C), making it a potentially useful transcriptional reporter for loss of sex-specific gene silencing.

### Loss of *stwl* results in derepression of testis-enriched genes in S2 cells and ovaries

Even when assayed mutant tissues appear morphologically similar to wild-type, the pleiotropic functions of Stwl nonetheless make it challenging to identify which genes are specifically misregulated as a consequence of Stwl loss. In order to further address this concern, we performed RNA-seq on a homogeneous tissue, using S2 cells treated with *stwl* dsRNA (see Methods). Immunoblotting against Stwl protein showed that *stwl* dsRNA treatment reduced Stwl protein levels by at least 80%, and RNA-Seq confirmed that *stwl* transcript was reduced by ∼85% (Fig 3A-B).

**Figure 3.**
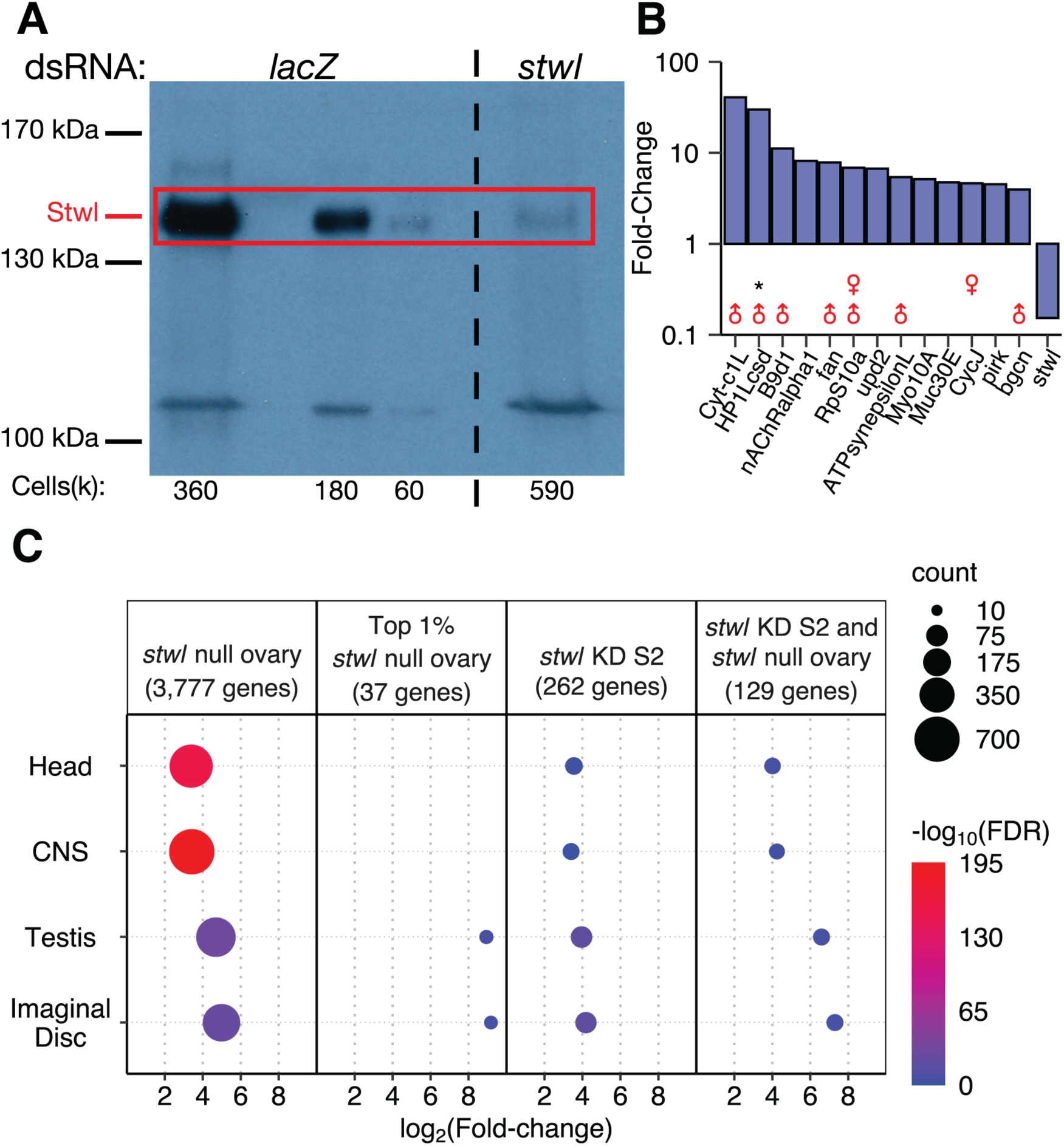
RNAi knockdown of *stwl* in S2 cells results in derepression of testis-enriched genes. (A) Western blot with anti-Stwl antibody of S2 cells treated with either control dsRNA (*lacZ*) or dsRNA targeting the *stwl* transcript. The estimated number of cells per lane (multiplied by 1000) is shown below the blot. (B) Fold-change of the 13 most affected annotated genes in *stwl* KD S2 cells relative to *lacZ* control. Male and female symbols indicate whether gene is testis- or ovary-enriched in wild-type. “*” marks genes that are part of the 59C4-59D testis-specific cluster. (C) FDR, count, and mean log_2_Fold-change (LFC) is plotted for each set of tissue-enriched genes that is overrepresented among ectopically expressed genes in *stwl* null ovaries and *stwl* dsRNA-treated S2 cells. Overrepresentation tests were also performed on the top 1% by LFC of ectopic genes in *stwl* null ovary, and of genes ectopic to both *stwl* null ovary and *stwl* dsRNA-treated S2 cells (for this intersect group, average LFC values in *stwl* null ovary are plotted). Only gene sets with FDR <0.05 are plotted.

Relative to loss of *stwl* in ovaries, *stwl* dsRNA treatment of S2 cells had a more subtle effect on transcript abundance and little effect on TEs (Supplementary Fig 11). Many fewer genes are expressed in S2 cells, which are male and hematopoietic-derived (Schneider, 1972; Zhang et al., 2010).

We found though that testis-enriched genes are among the most highly upregulated genes in *stwl* dsRNA-treated S2 cells, including a member of the 59C4-59D cluster (Fig 3B). Due to the very low average transcript abundance at this cluster in S2 cells, most of these genes were removed from the differential expression analysis performed by DESeq2 (Love et al., 2014) (Supplementary Table 5). Our further analysis of ectopically expressed genes, which was not limited by low average counts, found that 5/12 of the 59C4-59D genes upregulated in *stwl* null ovary are ectopically expressed in *stwl* dsRNA-treated S2 cells (Supplementary Table 4). Testis-enriched genes are over-represented among the top 1% of ectopically expressed genes in *stwl* dsRNA-treated S2 cells (Fig 2C).

For comparison, we applied the same methodology to our ovary data and found that 3,777 genes are ectopically expressed in *stwl* null ovaries. Head- and CNS-enriched genes are highly overrepresented among these ectopically enriched genes, as are testis- and imaginal-disc enriched genes, though to a lesser degree (Fig 3C). However, testis- and imaginal disc-enriched genes are the only upregulated tissue classes among the top 1% of ectopically expressed genes by LFC (Fig 3C). We found that 49% (129/262) of ectopically expressed genes in *stwl*-dsRNA treated S2 cells are also ectopically expressed in *stwl* null ovaries. This subset of genes is highly enriched for testis, imaginal disc, head, and CNS transcripts (Fig 3C). We conclude that *stwl* functions to repress genes with male-enriched expression, including in somatic tissue culture cells and ovaries.

### Stwl regulates key sex-determination and differentiation transcripts

Similar phenotypes of ectopic expression and upregulation of non-ovarian genes in female gonads have been reported for the female sex-determination gene *Sxl*, the H3K9me3 pathway members *egg*, *wde*, and *hp1a*, and the differentiation factor *bam* (Salz et al., 2017; Shapiro-Kulnane et al., 2015; Smolko et al., 2018). *Sxl* is required in ovaries for female sex determination; one of its critical functions is silencing (via the H3K9me3 pathway) of the male germline determining protein PHF7, a histone reader whose expression is necessary and sufficient for induction of spermatogenesis in the germline (Yang et al., 2012, 2017). PHF7 induction in female germ cells is also necessary for induction of the tumorous germ cell phenotype of *Sxl* defective ovaries (Shapiro-Kulnane et al., 2015). In wild-type female germ cells, transcription of *Phf7* is initiated from a TSS in the second exon, which results in truncation of the 5’ UTR of the female-specific transcript and absence of Phf7 protein in ovaries (Fig 4A). We found that the male-specific 5’ UTR is consistently and ectopically expressed in *stwl* null ovaries, regardless of age, but not in *stwl*-dsRNA treated S2 cells. Therefore, *stwl* is required for silencing of male-specific programming in ovaries.

**Figure 4.**
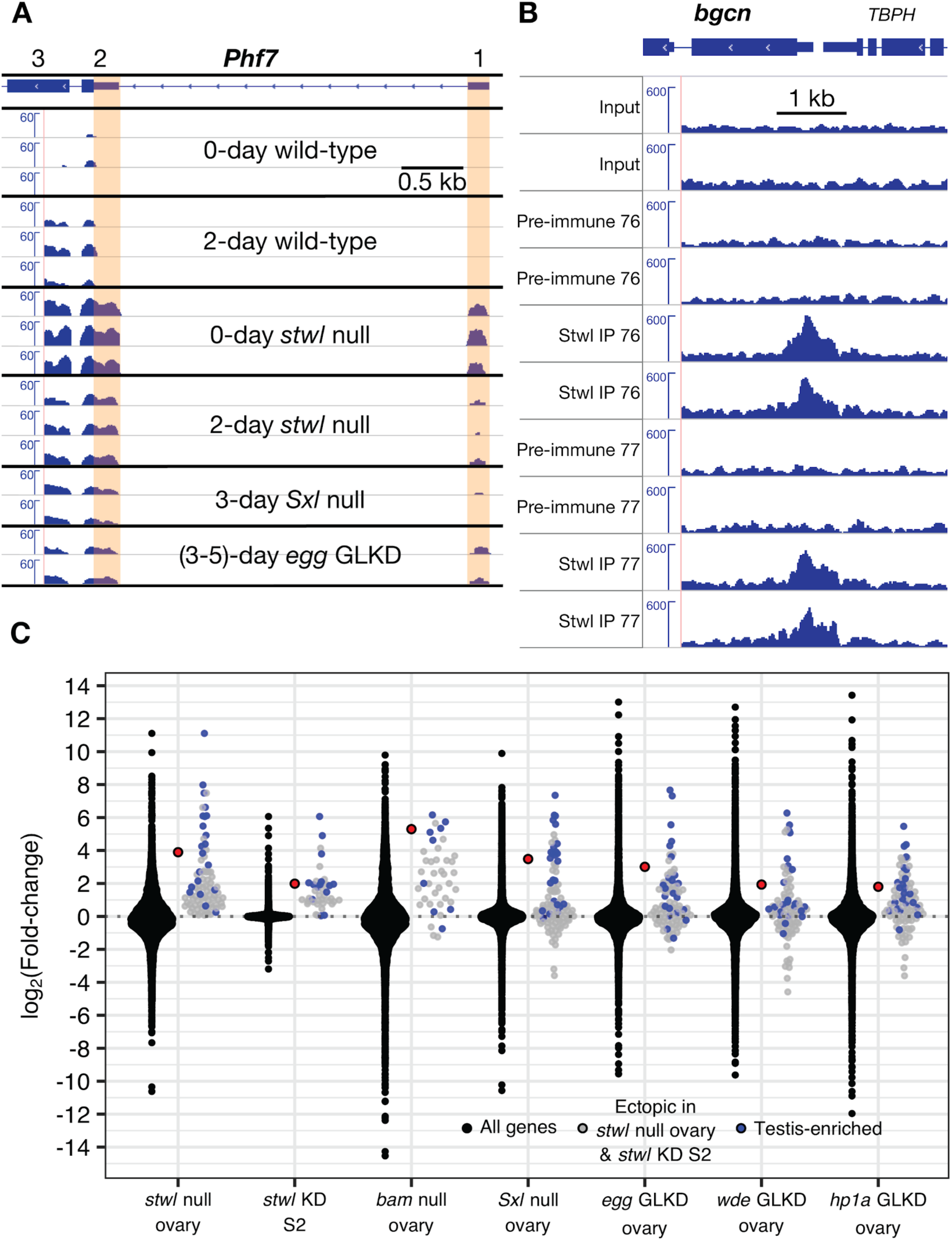
*Phf7* and *bgcn* are regulated by Stwl. (A) The male-specific 5’ UTR of *Phf7* (indicated by orange shading) is expressed in *stwl* deficient ovaries and S2 cells, as well as ovaries lacking *Sxl* and its downstream targets *egg*, *wde* (not shown), and *hp1a* (not shown) (Smolko et al., 2018). Reads were normalized to 1x depth of coverage and visualized in IGV, with *Phf7* shown in 3’ to 5’ orientation. Exons 1, 2, and 3 are indicated; exon 1 and part of exon 2 are male-specific. (B) *bgcn* is bound by Stwl in ChIP-Seq with two different anti-Stwl antibodies (76 and 77). Two independent replicates of each condition are shown. Reads were corrected for GC-bias, scaled to RPKM, and visualized in IGV. (C) Genes ectopically expressed in both *stwl* null ovary and *stwl*-dsRNA treated S2 cells (including *bgcn*, in red) are also upregulated in GSC mutants.

Since the overlap of genes ectopically expressed in *stwl* deficient ovaries and S2 cells is so striking, we predicted that upregulation of these genes may be consistent among ovaries exhibiting germline defects. Indeed, we found that these genes are generally upregulated in *bam, piwi,* and *Sxl* null ovaries, as well as *egg, wde,* and *hp1a* GLKD ovaries (Fig 4C). Of note, we find that *bgcn* transcripts are highly upregulated in each of the null and GLKD ovary datasets we examined. Bgcn binds to Bam and suppresses mRNAs associated with germline stem cell renewal, and both proteins are required to promote differentiation of developing cystoblasts.

In order to determine whether the effects of Stwl on gene expression that we discovered are direct or indirect, we developed two ChIP-grade antibodies against the protein and performed ChIP-Seq experiments on S2 cells. Both Stwl antibodies met our antibody validation criteria, including recognition of the target protein in immunoblotting and immunofluorescence experiments, successful immunoprecipitation of the target protein, and low background in immunoblots of S2 cells (Supplementary Figs 1-3, 12, Fig 3A). Our ChIP-Seq experiments produced a robust set of peaks when compared to both input and mock samples (Supplementary Fig 5).

We found very strong fold-enrichment of Stwl at the *bgcn* promoter (4.1-fold over input, IDR=1.3e^-5^) (Fig 4B). The peak at this locus is among the top 1% of Stwl peaks when ranked according to fold-enrichment. *bgcn* transcript is expressed at very low levels in *Drosophila* ovaries; its expression is limited largely to GSCs, where it is critical for promoting asymmetrical division into cystoblast daughters (Ohlstein et al., 2000). Loss of *bgcn* results in a tumorous ovary phenotype, as GSCs proliferate without differentiating into cystoblast daughters. While overexpression of Bam, the binding partner of Bgcn, results in GSC maintenance defects, this defect is not observed when Bgcn overexpression is driven in early germ cells (McCarthy et al., 2018; Ohlstein et al., 2000). We conclude that Stwl directly regulates expression of *bgcn* in the ovary and posit that aberrant expression of *bgcn* caused by Stwl loss results in activation of male-specific programming in the female germline.

### Stwl binding peak profiles are similar to known insulator binding proteins

We annotated 2,143 Stwl binding sites across the genome using ChIPseeker (Yu et al., 2015). Stwl is highly enriched at promoters, centered ∼150 bp upstream of transcription start sites (Fig 5A). To understand this pattern more deeply we compared our Stwl ChIP-Seq profile to the ModERN consortium data of 475 ChIP-Seq experiments on Gfp-tagged DNA and chromatin binding proteins in *D. melanogaster* embryos and larvae (Kudron et al., 2018) using the Genomic Association Tester (GAT) program (Heger et al., 2013). Briefly, GAT simulates a null distribution of peaks based on the size of each peakset, then estimates the number of overlaps expected by chance and compares this to the number of observed overlaps. We examined the most similar binding profiles according to fold-enrichment and % overlap. Reassuringly, ChIP-Seq against Stwl-GFP from ModERN had the most similar binding profile to our Stwl peakset, according to fold-enrichment (Supplementary Table 6). We also found that Stwl ChIP-Seq profiles were highly similar to a number of established and putative insulator binding proteins, including BEAF-32, CTCF, Su(Hw), Hmr, and Lhr (Fig 5B).

**Figure 5.**
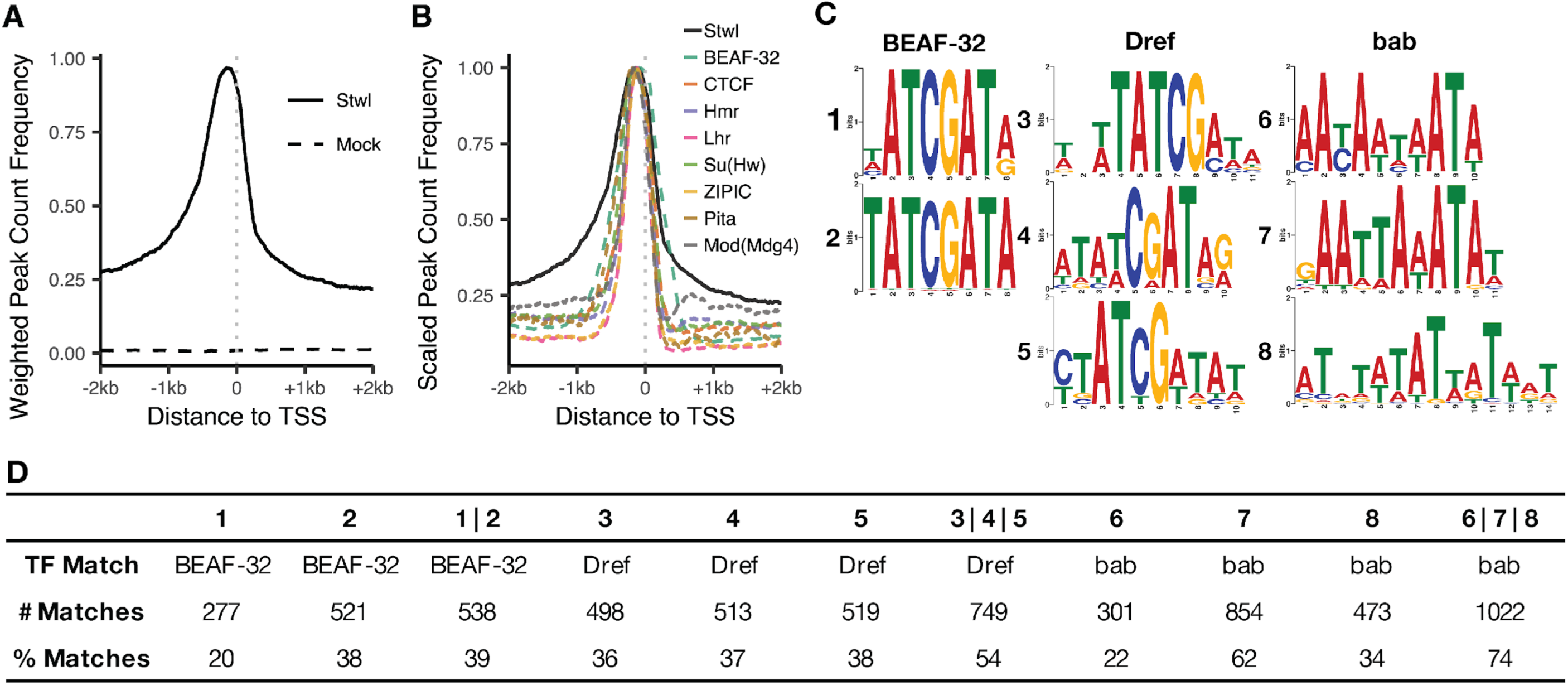
Stwl binding sites overlap with insulator-protein bindings sites. (A) Peak density of promoters bound by Stwl and mock antibodies. Frequency for each condition is weighted by the number of peaks present in the displayed 4 kb space. (B) Peak density of promoters bound by Stwl and known insulator binding proteins. Frequency for each protein is scaled to a maximum of 1. (C) Enriched motifs identified in narrow Stwl peaks using Meme Suite (Bailey et al., 2009). (D) For each motif from (C), we include the transcription factor that motif most closely matches with and the number of Stwl peaks that contain the given motif (# Matches). The % Matches identify the percent of the given motifs found in 1379 narrow Stwl peaks. Union columns (for example, 1|2) describe the number and % of narrow Stwl peaks that match to 1 or more of the indicated motifs.

We utilized the Meme Suite to identify enriched binding motifs in S2 cell Stwl ChIP-Seq (Bailey et al., 2009). We found that Stwl peaks are enriched for DNA sequence motifs that are common to BEAF-32, Dref, and bab (Fig 5c). Dref is an insulator binding protein that is additionally required for telomere maintenance (Tue et al., 2017). bab, which we identified as ectopically expressed in *stwl* null and *stwl*-dsRNA-treated cells, plays an important role in female sex differentiation (Williams et al., 2008). The occurrence of insulator motifs in Stwl ChIP-Seq combined with the above binding profiles provides strong evidence that Stwl binds to insulators.

### Stwl localizes to repetitive DNA, including telomeric repeats, chromosome 4, and pericentromeric heterochromatin

Previous studies have shown that Stwl is required for heterochromatin maintenance and colocalizes with HP1 at heterochromatin-like structures at the nuclear periphery (Maines et al., 2007; Yi et al., 2009). We found that Stwl is highly enriched across the dot chromosome (chromosome 4), which is highly repetitive and mostly heterochromatic (Riddle and Elgin, 2018) (Fig 6A). Stwl is also enriched at pericentromeric heterochromatin on chromosome 2, especially at the heterochromatin-euchromatin boundary. Finally, we saw a marked increase in coverage at cytological region 31 on chromosome 2L. Each of these regions is also enriched in Hmr ChIP-Seq in S2 cells (Gerland et al., 2017). Hmr localization at chromosome 2 has also been identified via immunofluorescence (Thomae et al., 2013).

**Figure 6.**
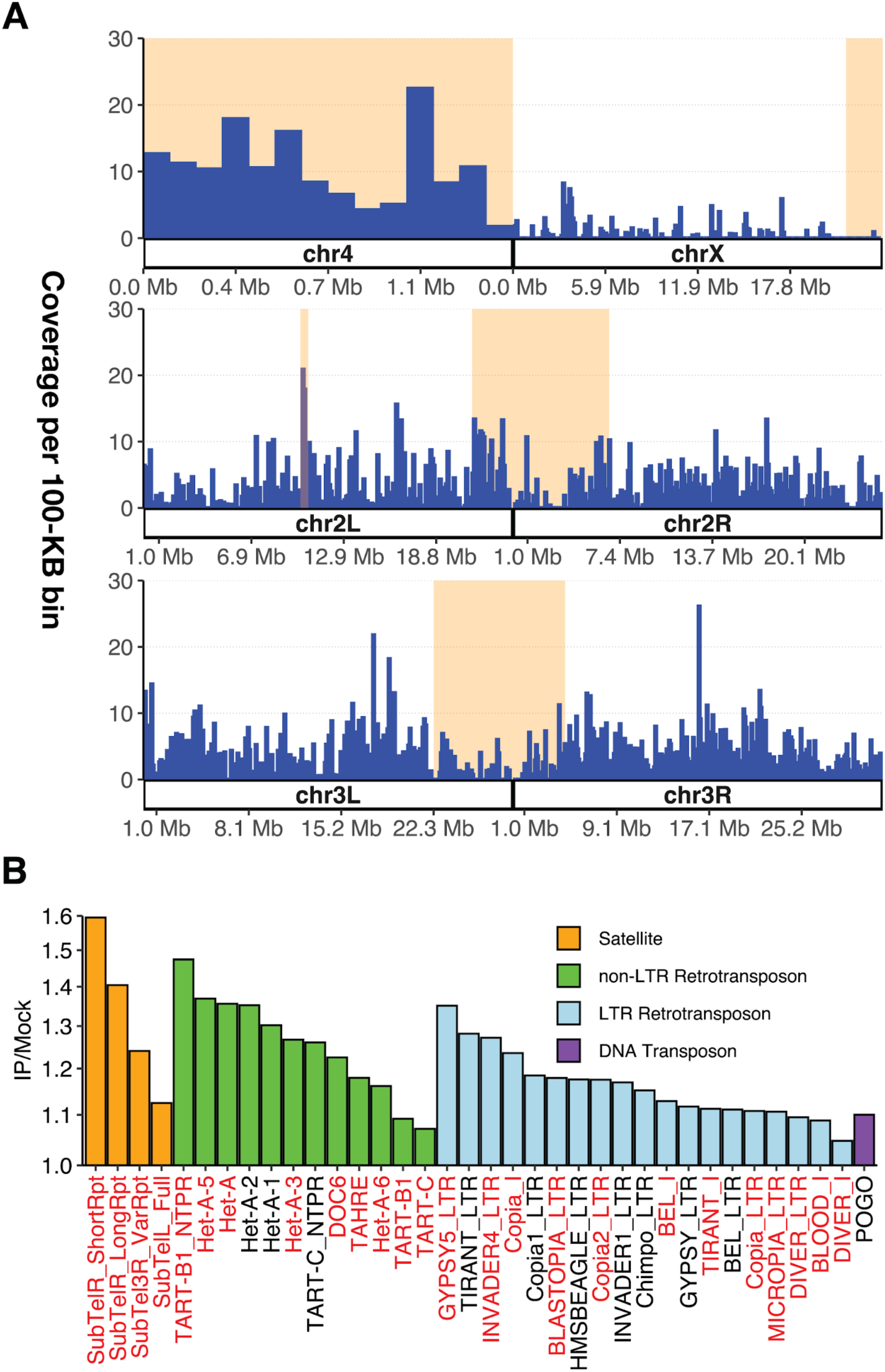
Stwl binds to repetitive DNA. (A) Percent (%) coverage of Stwl peaks (Y axis) per 100-kb bins of the genome (X axis). Shaded orange areas represent constitutive heterochromatin (pericentromeric regions and chromosome 4). Cytological region 31 on chromosome 2L is also highlighted. (B) Fold-change of read count abundance of repetitive elements for IP/mock comparison. Y-axis is in log2-scale. All significantly enriched elements (adj p <.01) are plotted. Red elements are upregulated in *stwl* null ovaries. All of the satellite and non-LTR retrotransposon sequences (except DOC6) are telomeric.

We next asked whether repetitive DNA, including satellite and transposable element sequences, are enriched among Stwl peaks. Since peak calling methods are not robust to repetitive DNA, we re-analyzed our ChIP-Seq data and instead calculated differential enrichment of reads in IP samples relative to mock (see Methods). This differential enrichment analysis identified repetitive DNAs enriched in Stwl IP samples (Fig 6B). All of these repeats passed a FDR threshold of 0.05, but the fold-changes of significantly enriched repeats were all less than 2. We note, however, that peak-calling algorithms can robustly identify enriched regions of DNA where fold-change of IP/mock is very low. In our own peakset, Stwl-bound sites passed IDR thresholding and were replicated in both antibodies, despite fold-change values as low as 1.2; the median fold-change for enrichment among all Stwl-bound peaks was 2.0. We are therefore confident that our Stwl ChIP-Seq has identified binding to repetitive DNA.

We found that Stwl ChIP-Seq is enriched for LTR retrotransposons, specifically members of the *copia*, *gypsy*, and *bel* superfamilies (Fig 6B). *Copia* elements are among the most highly upregulated transcripts in *stwl* null ovaries, and Stwl was enriched on both the *Copia* LTR and internal sequences. However, for other LTR retrotransposons such as *Bel* and *gypsy* this enrichment only occurred along the LTR component and not the internal region. We note that we did not detect motifs indicative of *gypsy* insulator binding in our motif enrichment analysis. These results suggest that Stwl might be involved in regulating these retrotransposons via their LTR regions. Alternatively, Stwl may be binding to heterochromatic fragments rather than regulating full-length active elements.

We were surprised to find that telomere-associated sequences are consistently enriched in Stwl IP (Fig 6B). With the exception of the *Jockey* family element *Doc6*, all enriched satellites and non-LTR retrotransposons are telomeric. These include telomeric satellite sequences and each of the members of the telomeric HTT array, *Het-A*, *TAHRE*, and *TART*. Furthermore, we found that Stwl peaks are highly similar to peaks generated from ChIP-Seq against the transcription factor pzg, and that Stwl shares DNA-binding motifs with Dref (Supplementary Table 6, Fig 5C). Each of these factors localizes to and is necessary for telomere maintenance (Andreyeva et al., 2005; Silva-Sousa et al., 2013). Lastly, we found that a majority of Stwl-bound telomeric repeat sequences are also upregulated in *stwl* null ovaries (gene names in red in Fig 6B). These findings suggest that Stwl localizes to telomeres and represses expression of telomeric repeats.

## Discussion

Identifying the molecular functions of Stwl is especially challenging due to its pleiotropic activity, including in GSC maintenance, oocyte determination, DNA damage response, and TE repression. Inferring Stwl function is further complicated by the consequences of *stwl* loss, e.g. apoptosis and eventual loss of the female germline. The resulting alteration of cellular content could lead to the identification of misregulated transcripts in *stwl* mutants that do not correspond to actual targets of wild-type *stwl*. We sought to tease apart direct versus indirect effects when analyzing steady-state RNA profiles of tissues affected by *stwl* loss. First, we assayed ovaries at two stages of development, thereby incorporating ovarian developmental status as a factor in the generalized linear model for differential expression. Second, we looked at differential expression in S2 cells after *stwl* knockdown in order to assay Stwl function in a homogeneous tissue. By combining these two assays, we were able to identify genes that are consistently upregulated due to *stwl* loss.

### Stwl represses male-specific transcripts

We identified in *stwl* null ovaries a single cluster of highly upregulated, testis-enriched genes on chromosome 2R. Genes in this cluster are among the top 1% of upregulated genes in ovaries. The 59C4-59D cluster is located within a lamina-associated domain (LAD). Such structures are thought to specifically repress expression of testis-specific genes by tightly binding these gene clusters to the nuclear lamina and preventing their expression. With the exception of this cluster, we did not find an association between *stwl* loss and misregulation of testis-enriched gene clusters, or LADs. We also do not find that Stwl is binding to this region, or overlapping with LADs.

In addition to the misregulated 59C4-59D cluster, we found that testis-enriched genes show a mixed pattern of up- and downregulation in *stwl* null ovaries. However, these genes are consistently among the most upregulated, ectopically expressed genes in ovaries. Additionally, we found that *stwl* null ovaries express the male-specific transcript of the master sex determination factor *Phf7*. These misregulated genes are not directly bound by Stwl, suggesting that derepression of these transcripts may be a downstream consequence of *stwl* loss.

As with Stwl, pleiotropy in other GSC maintenance genes makes it difficult to disentangle molecular function from cellular requirement. Furthermore, large-scale changes in ovary composition create challenges for interpretation of data generated from ovaries deficient for GSC maintenance genes. In *bam* and *Sxl* null ovaries, GSC-like cells overproliferate to form tumor-like structures with stem-cell-like qualities and gene expression patterns (Chau et al., 2009; McKearin and Ohlstein, 1995). They also tend to express transcripts associated with early gametogenesis in both sexes, many of which are testis-enriched transcripts. One possible explanation for the perceived “masculinization” of the ovary as a result of *Sxl* or *bam* mutation is that it reflects an overabundance of transcripts expressed during the early stages of gametogenesis (Shapiro-Kulnane et al., 2015). This is not the case for *stwl*. First of all, *stwl* null ovaries exhibit a GSC loss phenotype, not an overproliferation of such cells. Second, and most convincingly, we find that *stwl*-dsRNA treated S2 cells ectopically express many of the same testis-enriched genes that we identified in *stwl* null ovaries. S2 cells are male-derived, but our analysis nonetheless finds that the affected genes are almost completely silent in the control cells.

We found that multiple genes within the 59C4-59D cluster are also derepressed in *stwl*-dsRNA treated S2 cells, as well as *Sxl* and *piwi* null ovaries, and *egg*, *wde*, and *hp1a* germline-knockdown ovaries. In each of these cases, many of the genes in the cluster are ectopically expressed relative to wild-type ovaries. We conclude that ectopic expression of the 59C4-59D cluster and other testis-enriched genes is a consistent reporter of the “masculinization” defect associated with *stwl*, *Sxl*, *bam*, *hp1a*, *wde* and *egg* mutants.

### Stwl regulates Bgcn

While male transcripts are upregulated and/or ectopically expressed in *stwl* mutants, our ChIP data suggests that Stwl does not bind at these loci, suggesting that the masculinization defect is an indirect consequence of *stwl* loss. One possibility is that these phenotypes are associated with ectopic expression of *bgcn*, which is typically restricted to GSCs and cystoblasts in ovaries, but widely and highly expressed throughout spermatogenesis (Insco et al., 2012; Ohlstein et al., 2000). *bgcn* is a strong candidate for Stwl regulation: its promoter is bound by Stwl, and it is highly upregulated in *stwl* null ovaries and *stwl* dsRNA-treated S2 cells. Furthermore, the expression of *bgcn* transcripts in ovaries is anti-correlated to Stwl, and females expressing a *hs-bgcn* transgene are sterile (Ohlstein et al., 2000). We propose that loss of Stwl in the developing germline cyst results in ectopic expression of Bgcn in these cells that eventually leads to female sterility (Fig 7).

**Figure 7.**
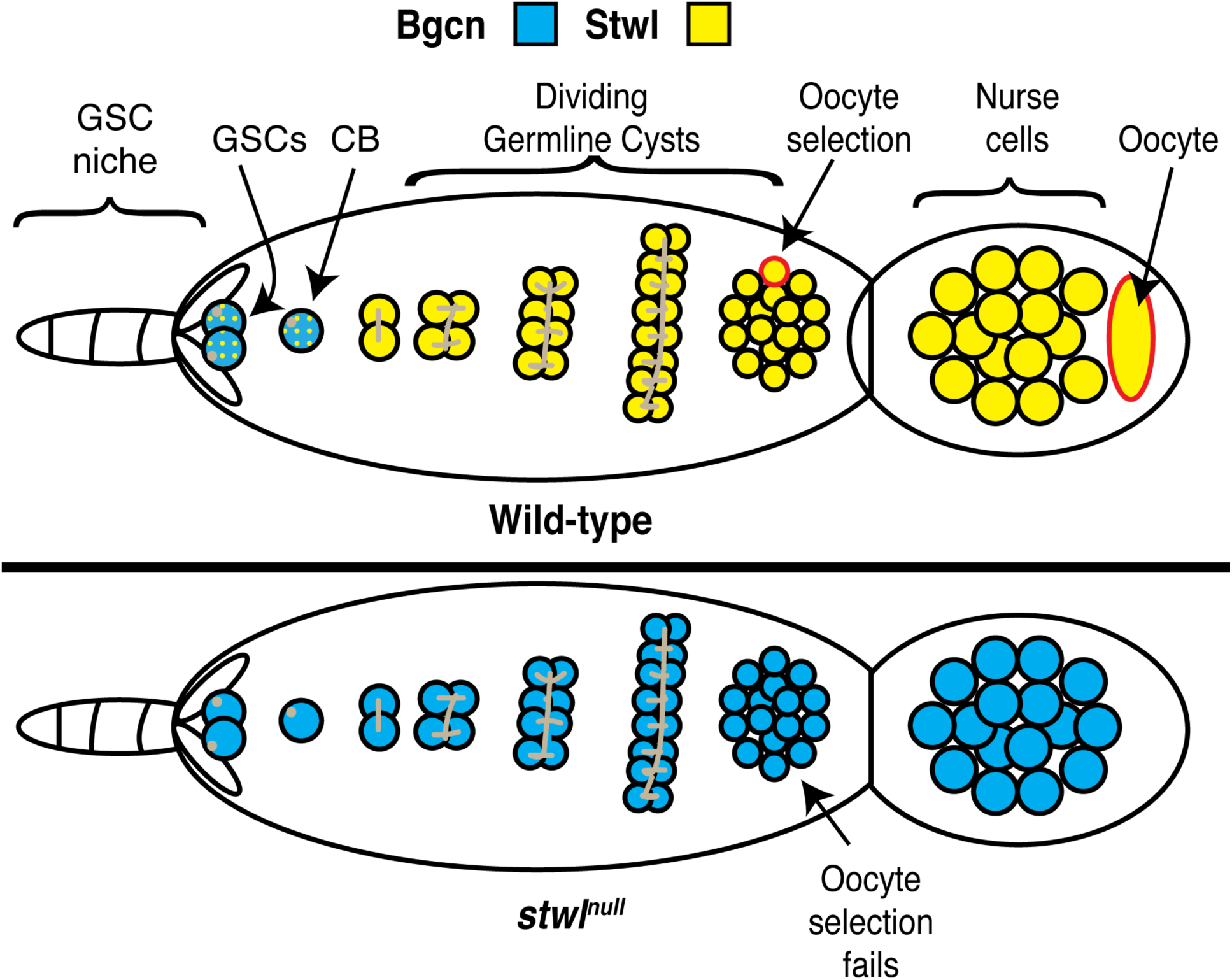
Model for Stwl as an essential regulator of Bgcn. In wild-type ovaries, Bgcn expression (blue) is restricted to GSCs and cystoblasts (marked by presence of round spectrosomes in tan). Stwl expression is low in these same cells and increases substantially in the developing germline cyst (with branched fusomes, in tan). Loss of Stwl results in an increase of Bgcn outside of GSCs and cystoblasts, causing expression of male transcripts that ultimately disrupts formation of the oocyte (red border) in the 16-cell cyst.

The molecular pathways in which Stwl functions to maintain oogenesis, either at the stage of germline stem cell retention or oocyte determination, may overlap significantly with pathways in which Bam and Bgcn are crucial actors. Stwl acts independently of and antagonistic to Bam: *bam* mutants present with GSC-tumorous ovarioles, while *stwl bam* double mutants form rudimentary germline cysts (Maines et al., 2007). Despite the fact that Bgcn has a defined and important role in the ovary, it is nonetheless largely silent throughout oogenesis. Overexpression of *bam* in GSCs results in loss of GSCs, similar to *stwl* nulls; surprisingly, *bgcn* overexpression does not result in GSC loss (Ohlstein et al., 2000). Nonetheless, non-specific expression of a *bgcn* transgene causes sterility in females, characterized by “small oocytes” in late stage egg chambers. We suspect that ectopic expression of *bgcn* outside of GSCs results in sterility. It is possible that one of Stwl’s functions in the female germline is to restrict expression of *bgcn* to GSCs. There is evidence that *stwl* is epistatic to *bgcn*: *bgcn stwl* double mutant ovaries form fusomes, which are not present in *bgcn* mutant ovaries (Park, 2007).

### Stwl accumulates at genomic insulators and heterochromatin

Loss of *stwl* results in derepression of repetitive elements, a phenotype that is also observed in *bam* and *piwi* mutant ovaries. While Piwi is a known regulator of TEs via the piRNA pathway, it is unclear whether upregulation of TEs in *bam* and *stwl* mutant ovaries reflects a direct role in TE silencing. In order to answer whether Stwl directly targets repetitive DNA and how it may be involved in TE control, we developed antibodies to Stwl and assayed Stwl binding in S2 cells. Our analyses indicate that Stwl binds to insulator elements. Most Stwl peaks are located just upstream of promoters; this binding profile is common among insulator-bound proteins. More directly, we identified strong sequence similarity between Stwl peaks and peaks from a number of insulator binding proteins, including BEAF-32, Dref, ZIPIC, Pita, Hmr and Su(Hw). In addition, we found that *stwl* peaks accumulate at heterochromatic loci, specifically pericentromeric heterochromatin boundaries, telomeres and the dot chromosome.

These data suggest that Stwl is an insulator binding protein, consistent with previous work showing that Stwl associates with insulator complexes in immunofluorescence experiments (Rohrbaugh et al., 2013). Insulators have multiple functions including blocking enhancer-promoter interactions and establishing boundaries to prevent the spread of chromatin modifications and to separate differentially expressed promoter pairs. Insulator-binding proteins, such as CP190, can also mediate long-range interactions (Saha et al., 2019; Vogelmann et al., 2014). If Stwl is involved in the formation of long-range interactions, it may promote tethering of euchromatic insulator sites to heterochromatic regions. We speculate that Stwl-bound sites are located adjacent to regions of repressed chromatin, and that loss of Stwl results in spreading of these repressed chromatin marks to neighboring loci. It is likely that Stwl performs its function as an insulator by establishing boundaries, in conjunction with other insulator-binding or heterochromatin-associated proteins, that ensure proper expression of nearby genes.

The *D. melanogaster* ovary is a complex mixture of cell types in the adult fly. Differentiated somatic cells function as support cells to shepherd germ cells towards their ultimate fate of producing viable gametes. Furthermore, each of these cellular lineages is derived from a small population of self-renewing stem cells. We suggest that insulators allow genes to have pleiotropic functions during development of complex tissues such as ovaries. Insulators add a layer of genomic complexity to gene regulation by disrupting enhancer-promoter interactions. The specific interplay of promoters, enhancers, and insulators during oogenesis is poorly understood but is likely key to explaining pleiotropic gene regulation in the developing ovary.

## Acknowledgements

We thank Chip Aquadro, Shuqing Ji, Heather Flores, Satyaki Prasad, Kevin Wei, Michael McGurk, and Dean Castillo for their input at various stages of this project. We also acknowledge Abdullah Ozer, Judhajeet Ray, Roman Spektor and Tawny Cuykendall for their invaluable guidance on ChIP-Seq library prep, Aaron Chen for his technical contributions, and Helen Salz for helpful conversations.

---

**Figure S1.**
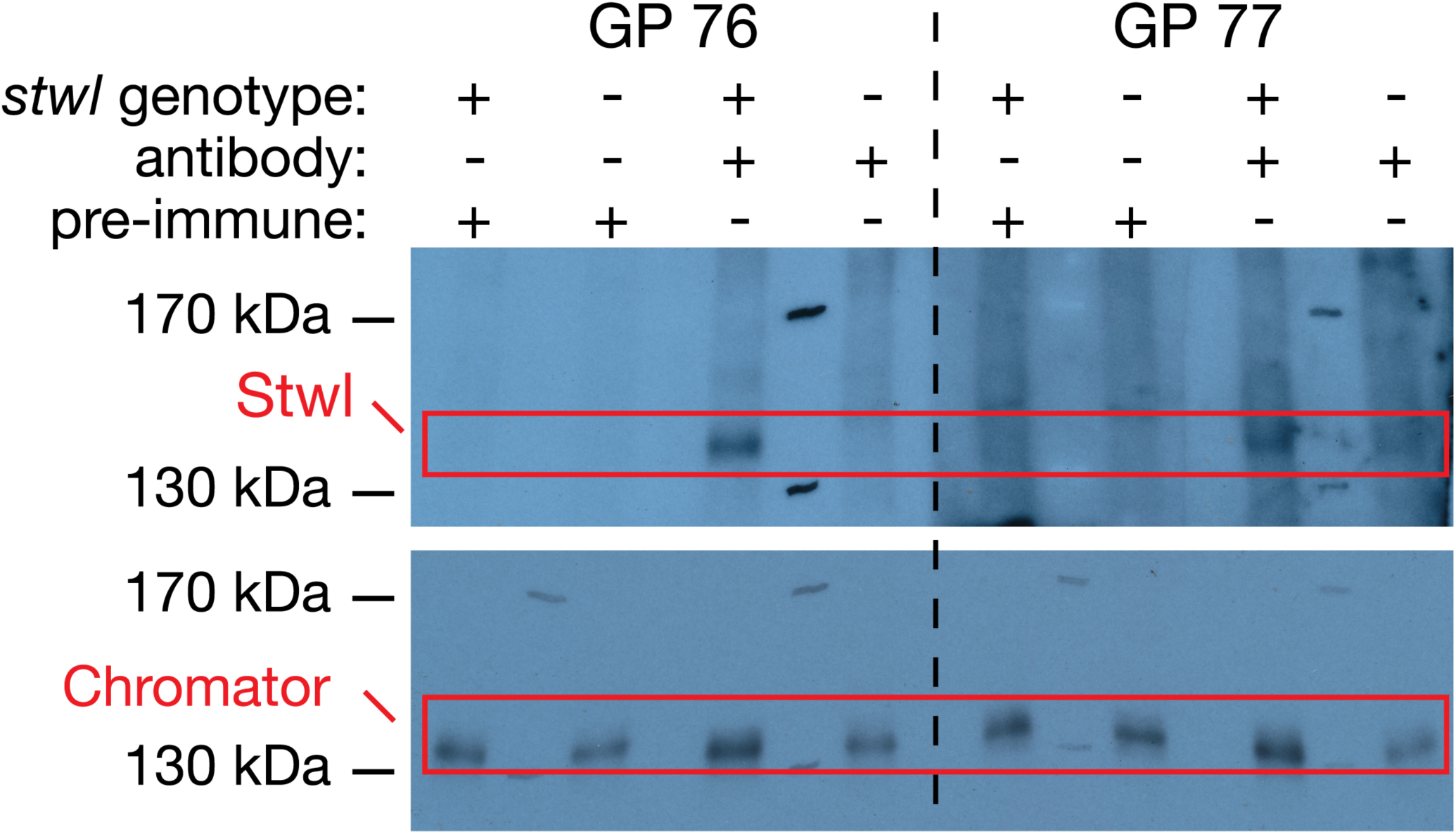
Stwl antibodies recognize a ∼130 kDA fragment. Western blots on whole-fly lysates from ∼10 *stwl^+^* (*y w* F10) and ∼10 *stwl* null (*stwl^j6c3^*/*Df(3L)Exel6122*) individuals aged 1-4 days. 6% SDS PAGE gel was loaded with lysates as indicated (row labelled “*stwl* genotype”), then transferred and probed with pre-immune or antibody sera of each animal, as indicated. The bottom panel shows the same membrane stripped and re-probed with a loading control (guinea pig ɑ-Chromator). Final-bleed serum of each Stwl antibody recognizes a ∼130 kDa fragment specific to *stwl^+^* lysates.

**Figure S2.**
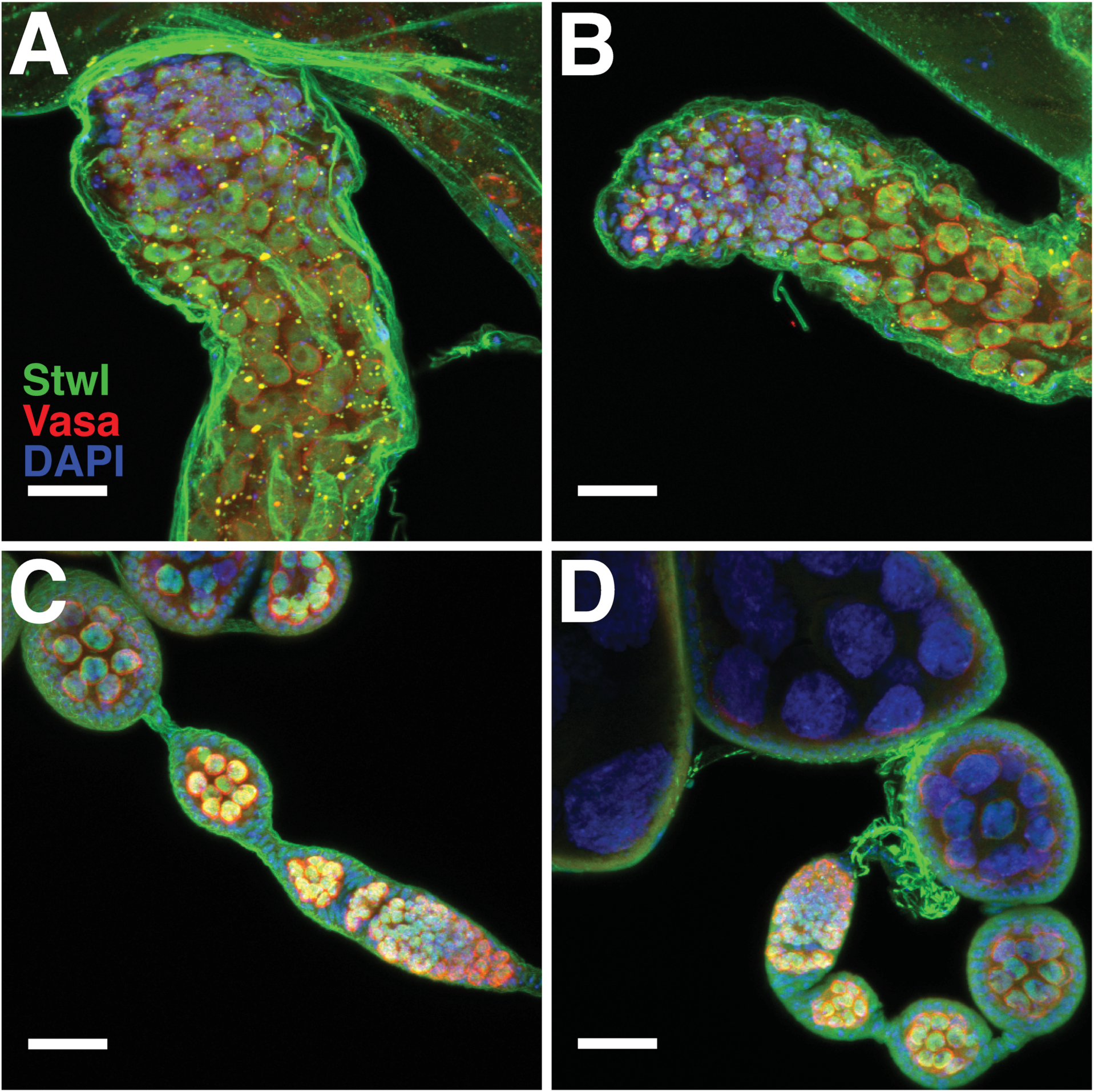
ɑ-Stwl sera label germ cell nuclei in *stwl^+^* testes (A, B) and ovaries (C, D). Tissues were dissected from *y w* F10 flies 10-15 days post-eclosion and immunostained with ɑ-Stwl sera from GP 76 (A, C) and GP 77 (B, D). Vasa labels germ cells, DAPI labels cell nuclei. All images are maximum-intensity projections from a z-series representing a depth of 10 µm. Scale bars are 20 µm.

**Figure S3.**
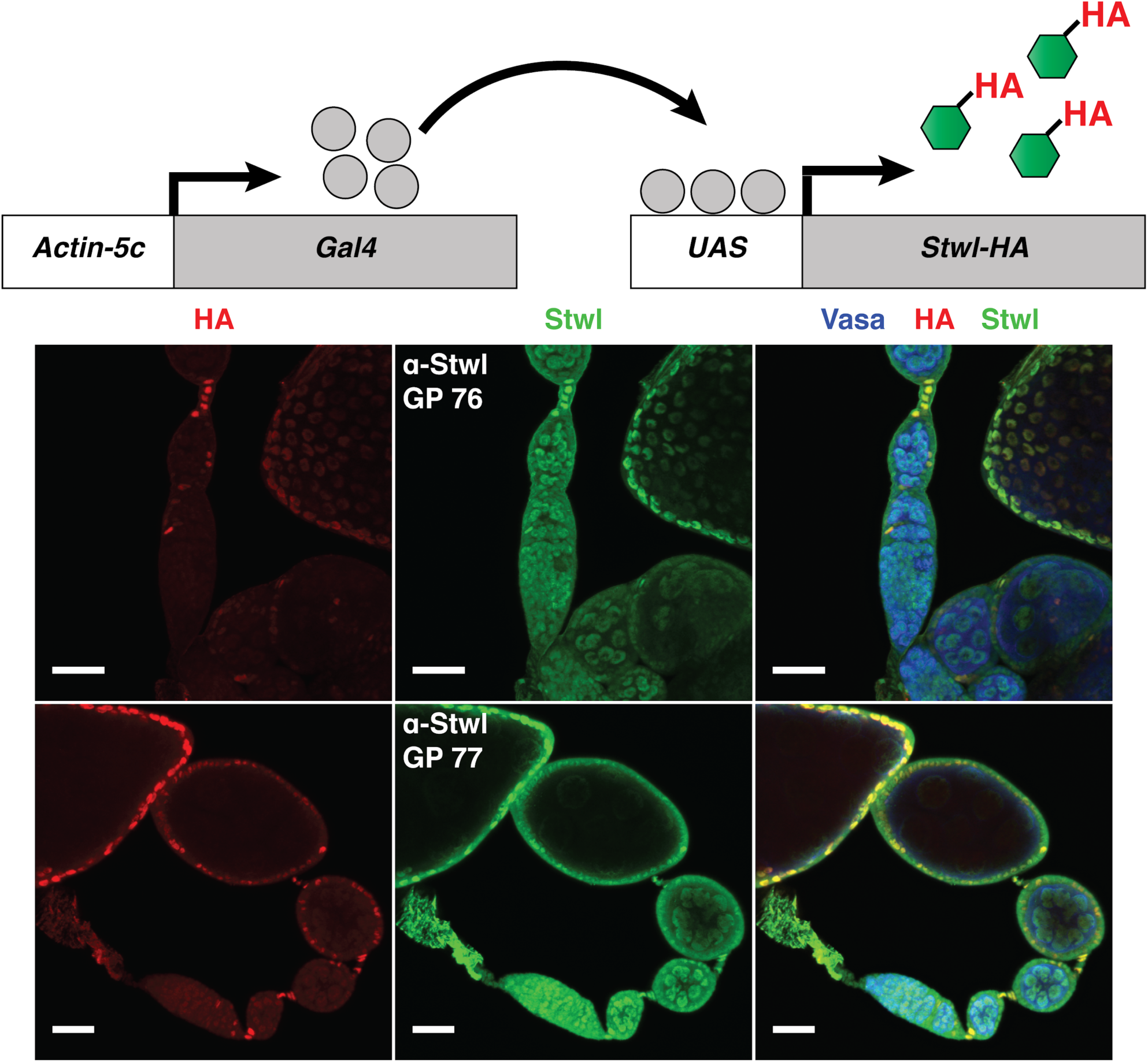
ɑ-Stwl sera detect ectopically expressed Stwl-HA protein. Ovaries were dissected from *Act5c-Gal4/UAS-stwl-HA* females 0-1 days post-eclosion. Ovaries were probed with ɑ-Vasa (germ cells), ɑ-HA, and either GP 76 or GP 77 ɑ-Stwl serum. HA signal recognizes cells in which Stwl-HA is being expressed; in these examples, expression is mostly limited to somatic cells (follicle cells and stalk cells). ɑ-Stwl signal for both antibodies clearly overlaps with HA signal, resulting in bright yellow foci in the composite image. All images are maximum-intensity projections from a z-series representing a depth of 10 µm. Scale bars are 20 µm.

**Figure S4.**
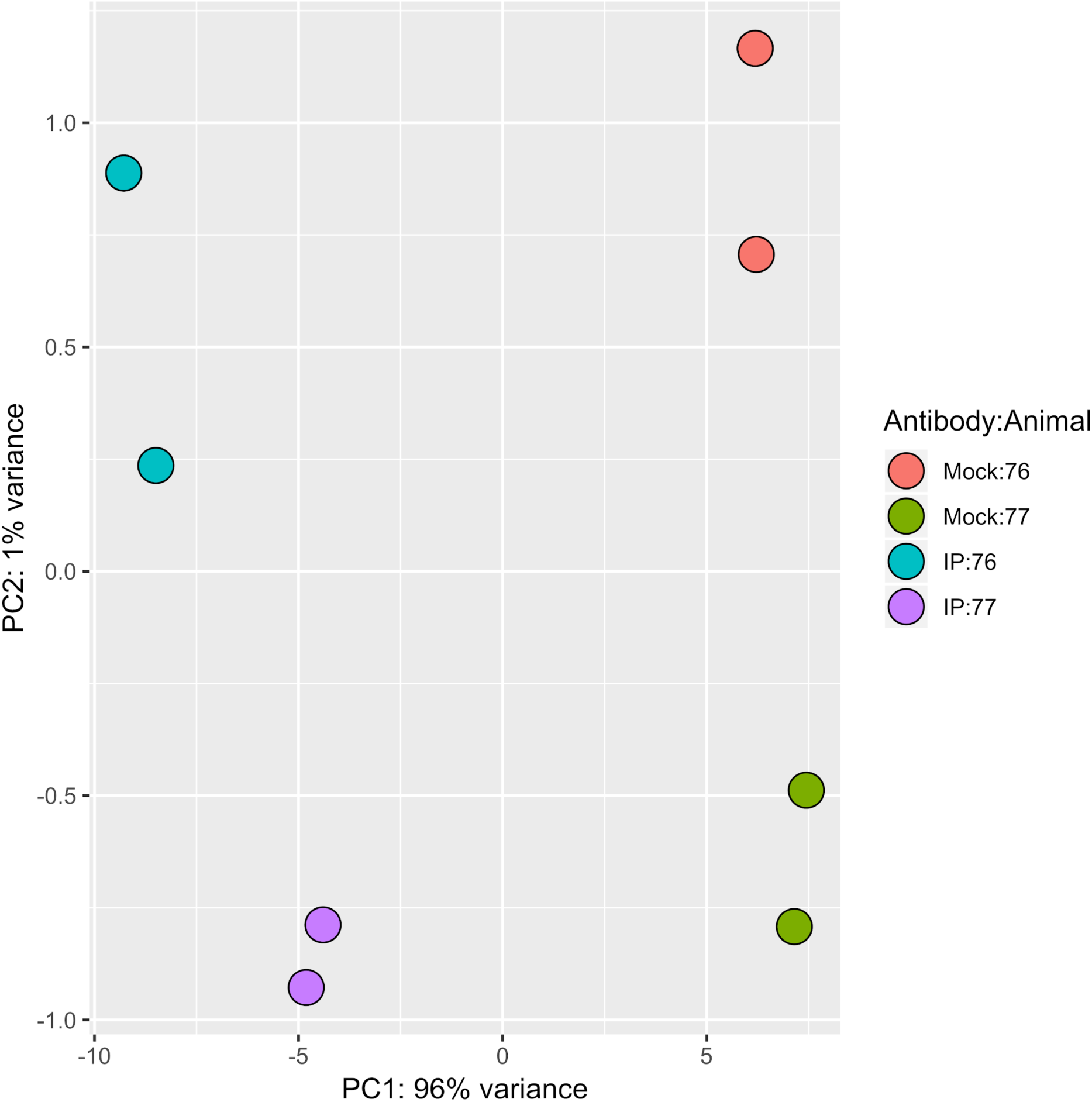
Principal components analysis (PCA) of ɑ-Stwl ChIP-Seq read counts separates mock from IP. PCA for read counts generated from alignment to genomic bins and repeat index. Experiments labeled as mock were performed with pre-immune sera, IPs were performed with Stwl antibodies. Antibodies were generated from two different animals (referred to as 76 and 77) using the same epitope. DNA was isolated from two pools of S2 cells (biological replicates). The majority of the variance in the data is contained in PC1 and is explained by differences between mock and IP condition, not by differences in the source animal or replicate pools.

**Figure S5.**
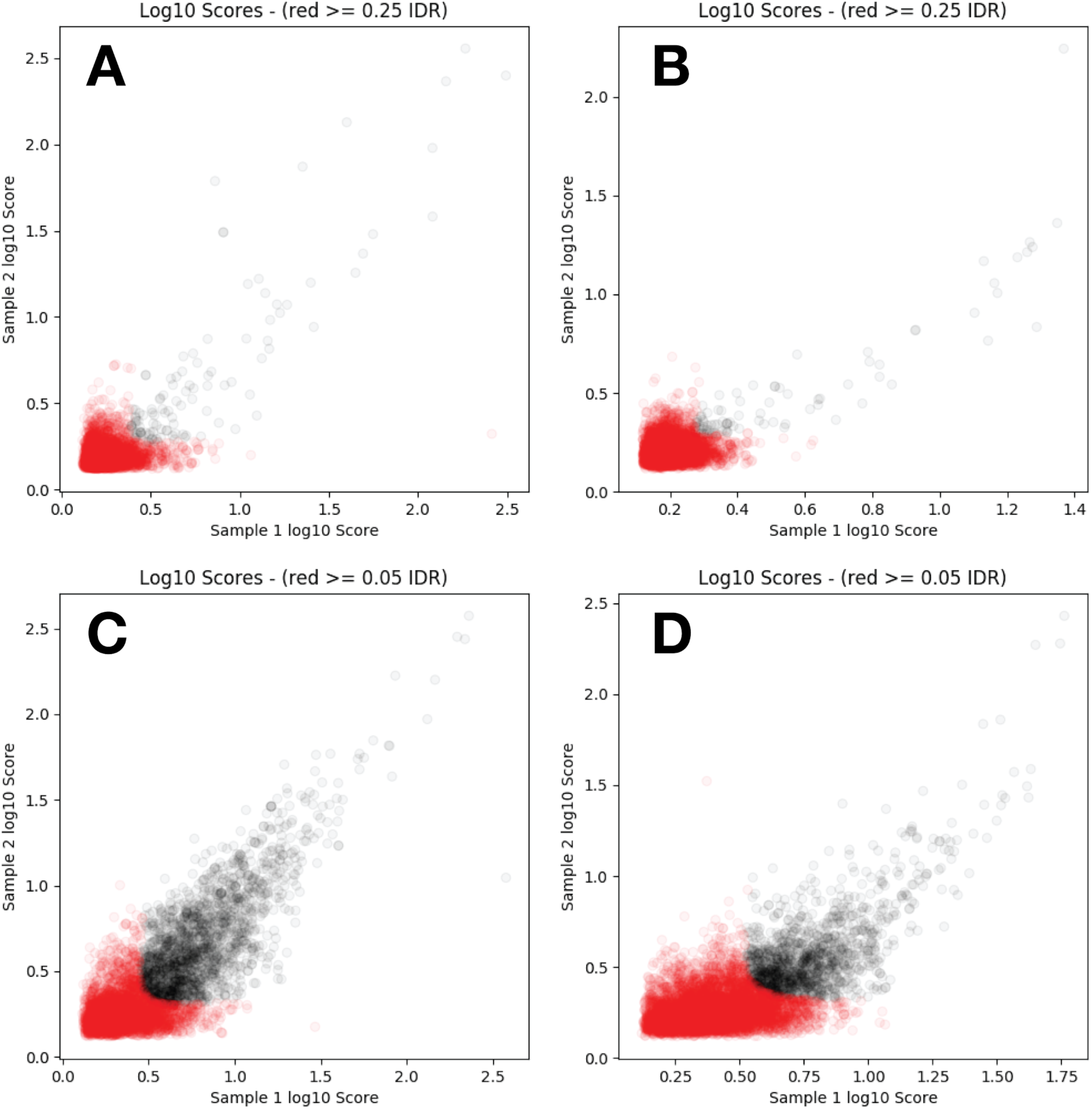
Stwl IP replicates create reproducible peaksets. IDR plots show the distribution of peak scores in replicate 1 (x-axis) vs replicate 2 (y-axis). Grey dots are reproducible peaks that pass the given IDR threshold, red dots are irreproducible peaks. Each dot represents a ChIP-Seq peak called in both replicates of a single antibody (C, D) or mock (A, B) experiment. Peak scores reflect the fold-enrichment of reads in the IP or mock sample relative to input. IDR identifies peaks whose signal intensities (i.e. scores) are similar in both replicates. Peaks with low signal intensity in both replicates do not pass the IDR threshold, but are useful for generating a background dataset for IDR analysis. Very few peaks were identified in mock ChIP-Seq experiments, even with a relaxed IDR threshold of 0.25.

**Figure S6.**
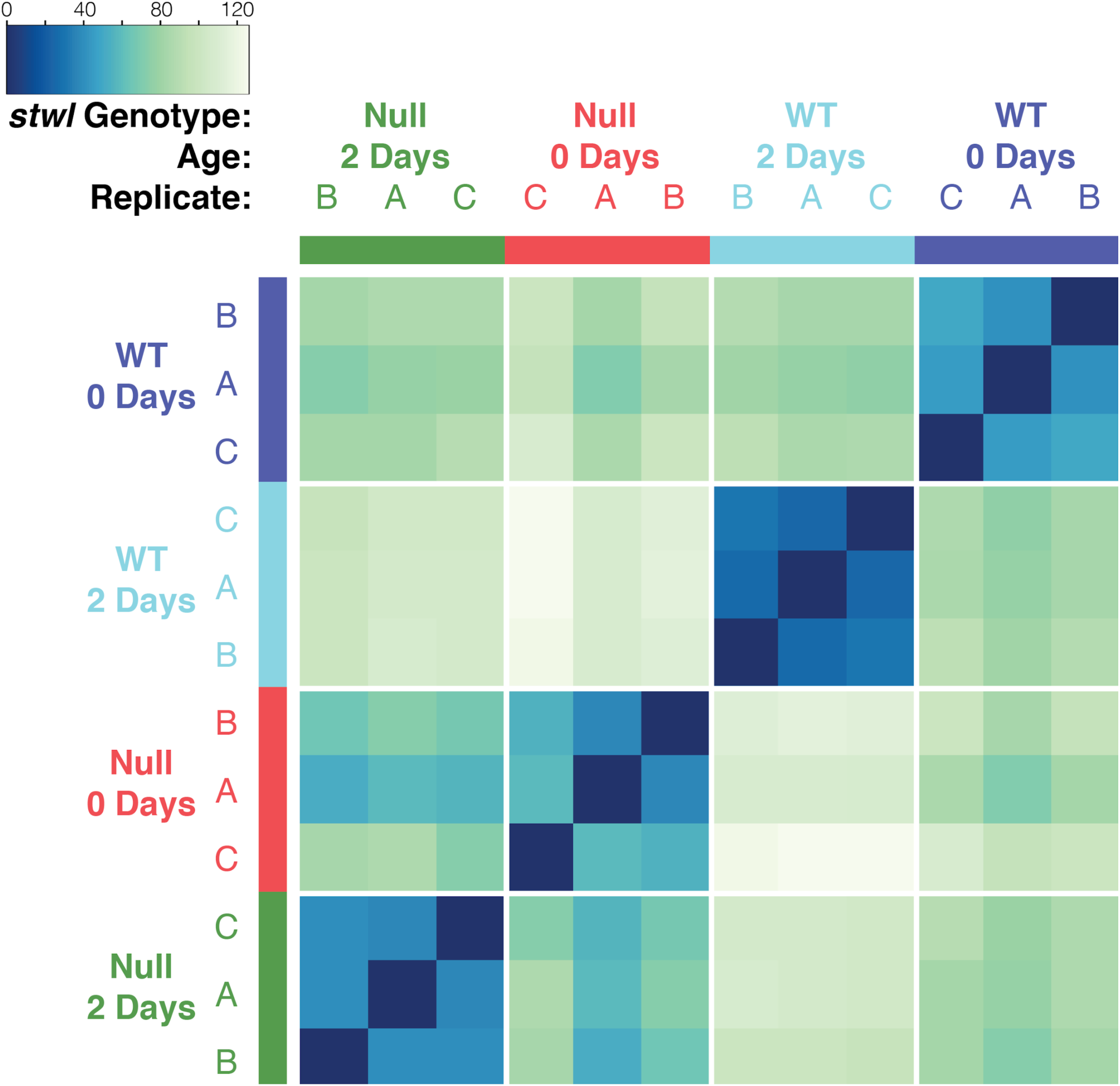
Sample-to-sample distance matrix of RNA-Seq samples. Read counts were regularized log transformed in DESeq, and the distance between samples was calculated based on these transformed count values. The heatmap is sorted by similarity after hierarchical clustering and color-coded according to distance, where dark blue cells indicate a distance of 0 (completely self-similar) and white cells a maximal distance (completely dissimilar). Samples within the same group (identical age and genotype) occur together and form blue clusters.

**Figure S7.**
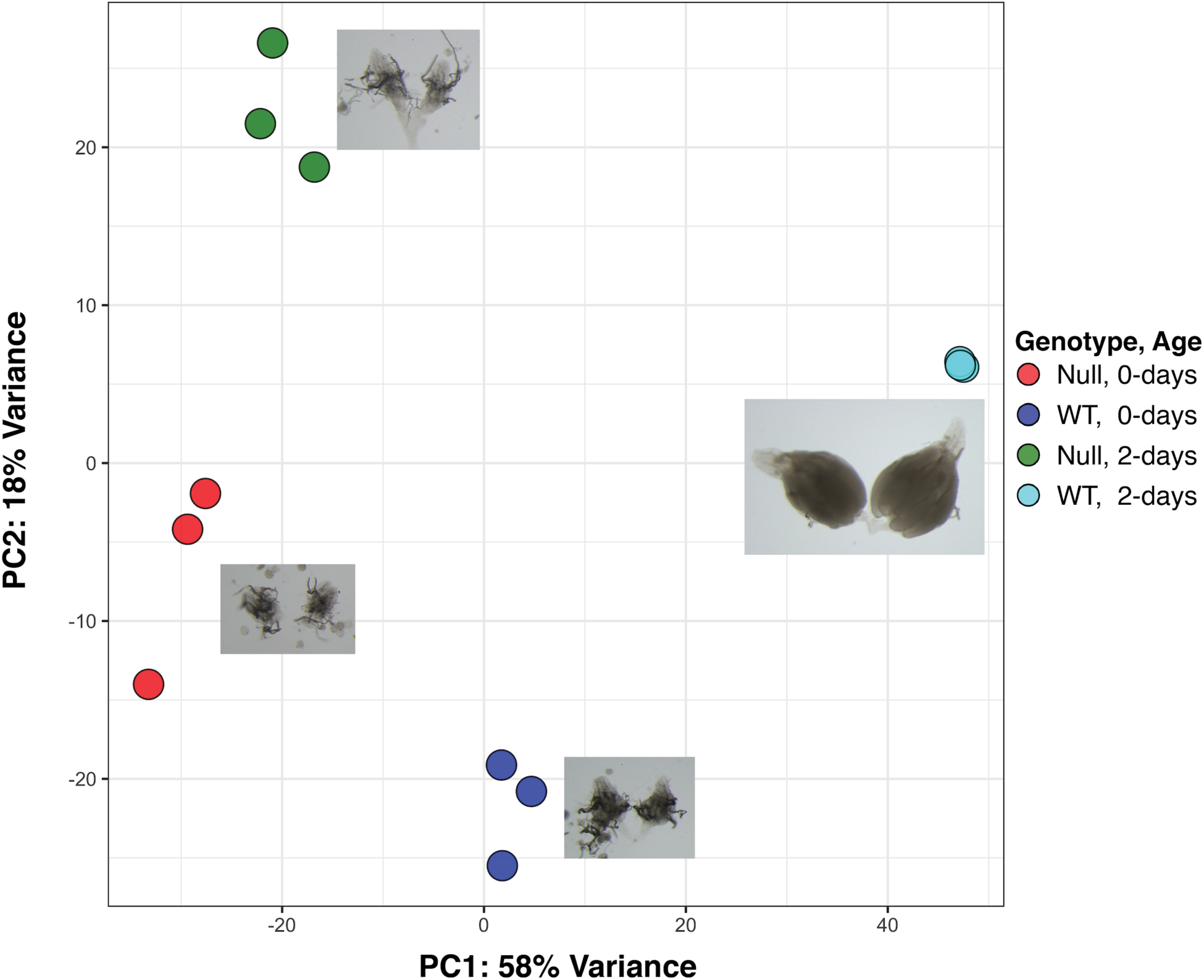
Principal Component Analysis (PCA) of RNA-Seq count matrices. PCA was performed on regularized log transformed read counts of the 500 most variable genes in the count matrix. Samples within the same group (identical age and genotype) cluster together, indicating minimal batch effects.

**Figure S8.**
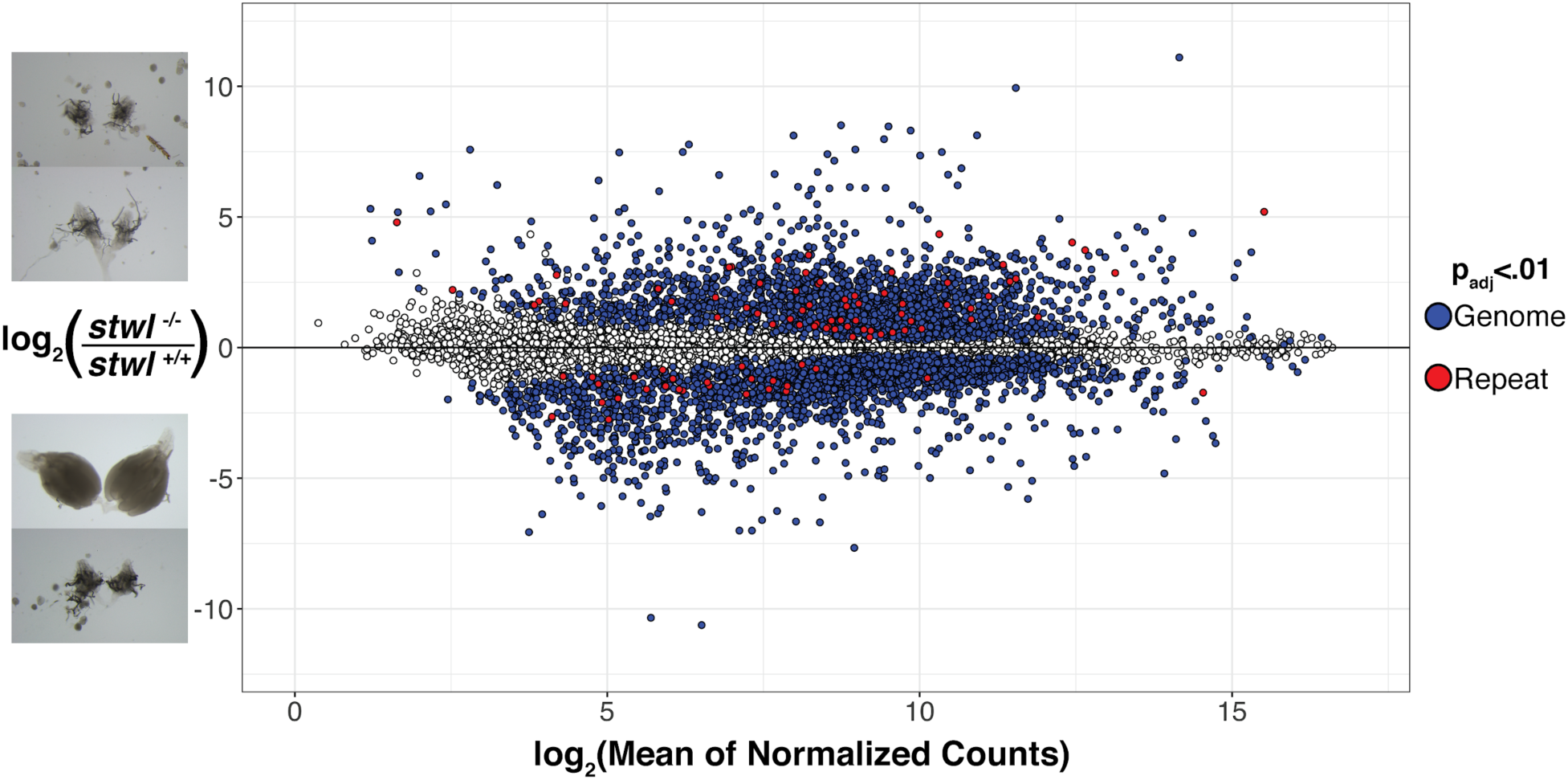
MA plot of RNA-Seq data from ovaries. Fold-change for each gene is plotted against its average transcript abundance across all assayed ovarian samples (wild-type and null). Transcript abundance is represented by counts normalized according to GC-content and library size. The log2(Fold-change) values (LFC) were “shrunk” to minimize the variance associated with low-count genes. Filled points (blue and red) identify genes which are differentially expressed (adjusted p-value <0.01) in this comparison. Red points represent entries from Repbase, blue points are from the genomic annotation.

**Figure S9.**
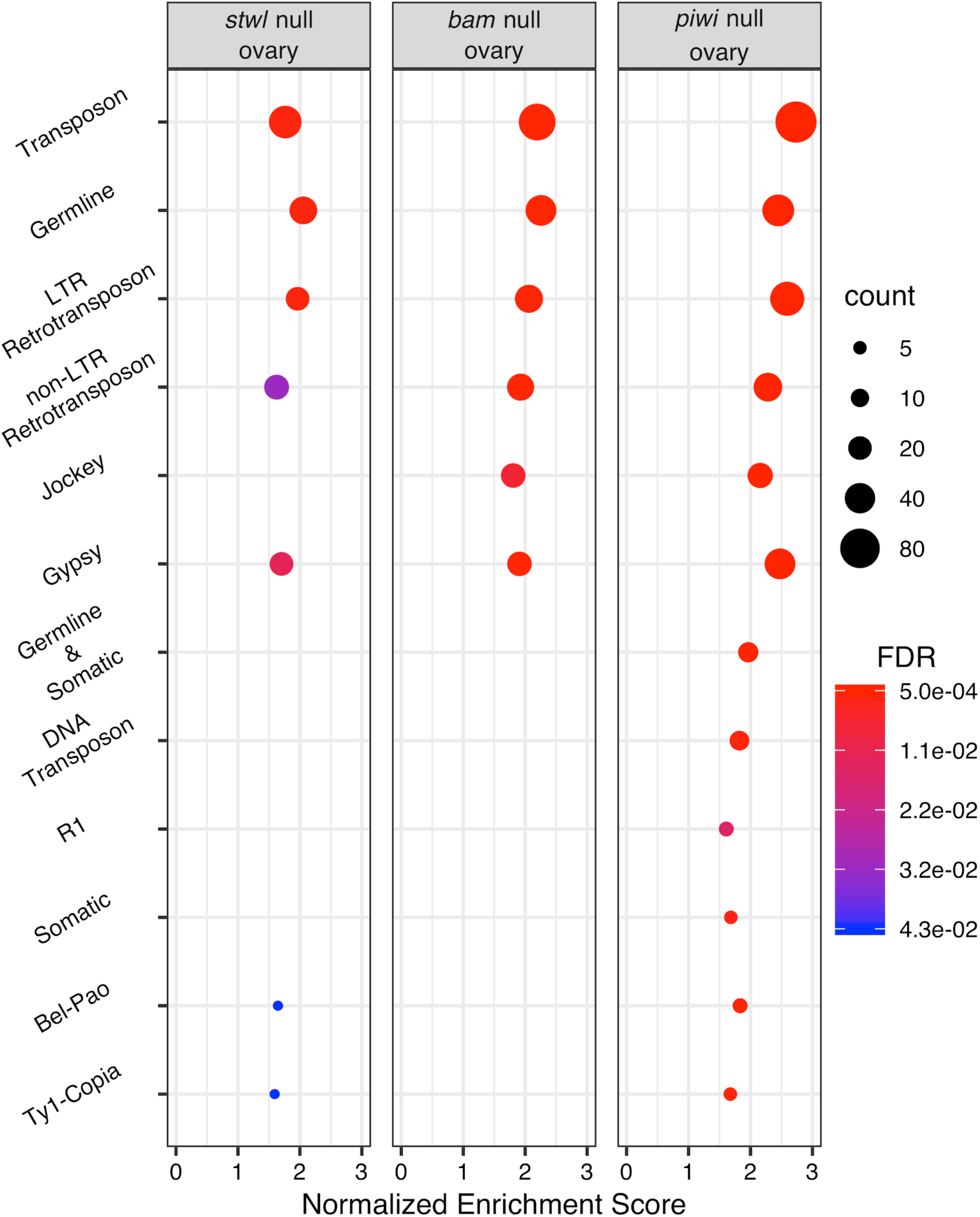
TE de-repression in *stwl*, *bam*, and *piwi* null ovaries. Comparison of Gene Set Enrichment Analysis (GSEA) results. Normalized Enrichment Score (NES) is plotted for each set of repetitive elements enriched among null/WT ovaries. Higher NES indicates that the gene set is more upregulated in null ovaries. Count represents the number of genes in that set. Only gene sets with FDR<0.05 are plotted. *Sxl* null ovaries were also analyzed but are not shown because they were not enriched for any repeat classes.

**Figure S10.**
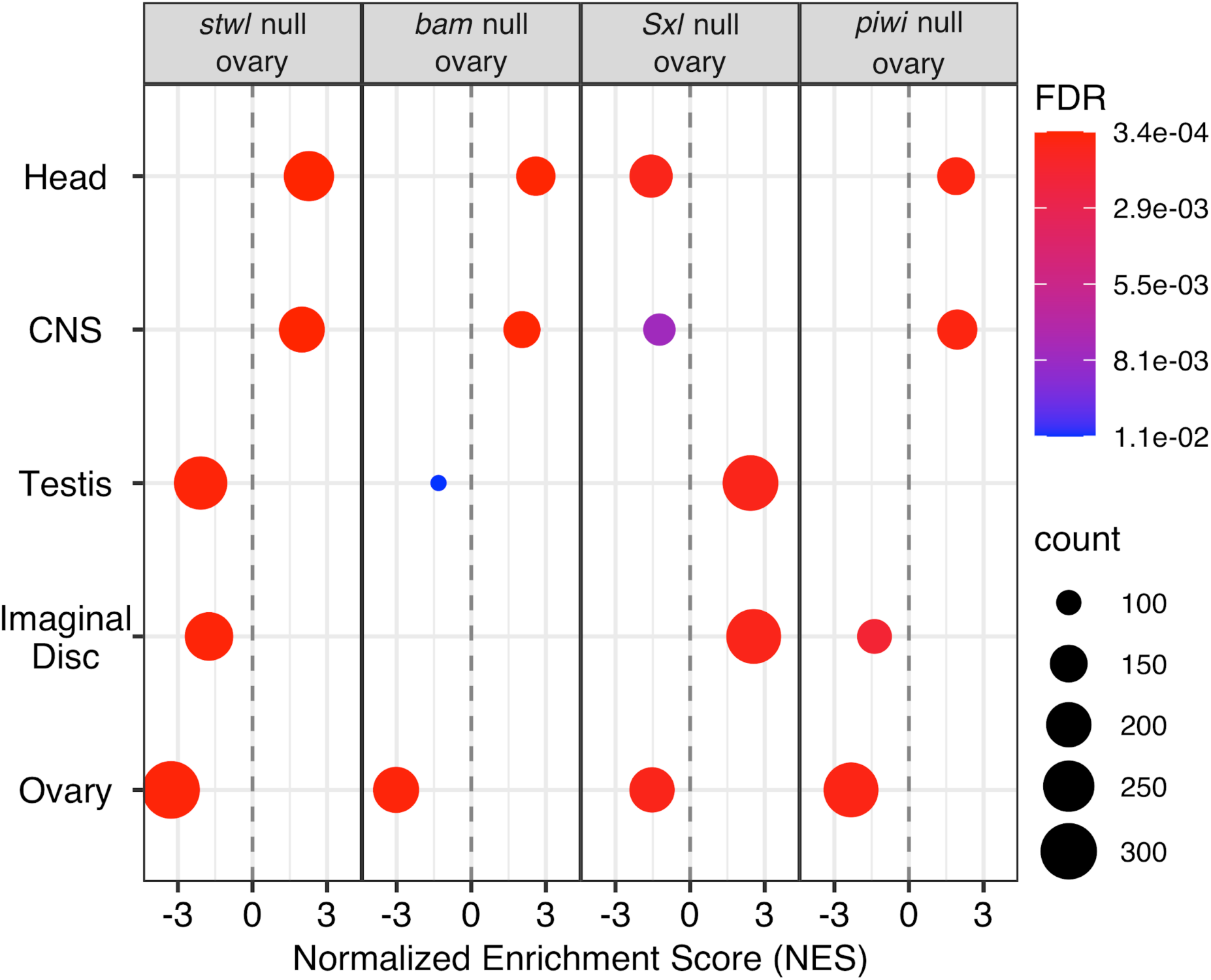
Comparison of Gene Set Enrichment Analysis (GSEA) results for tissue-enrichment. Normalized Enrichment Score (NES) is plotted for each set of tissue-enriched genes enriched among null/WT ovaries. Higher/lower NES indicates that the gene set is highly upregulated/downregulated in null ovaries. Count represents the number of genes in that set. Only gene sets with FDR<0.05 are plotted.

**Figure S11.**
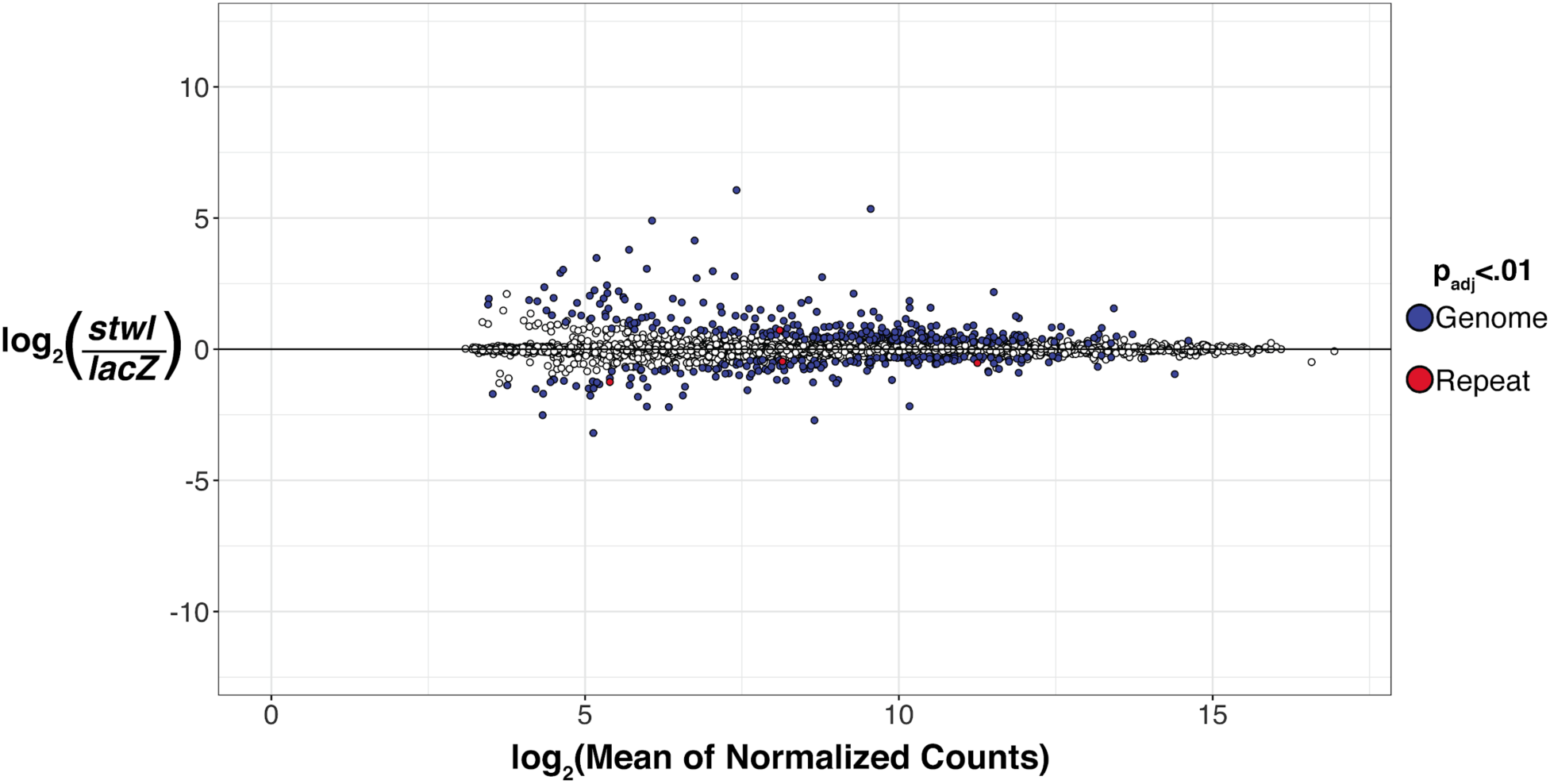
MA plot of RNA-Seq data from S2 cells. Fold-change for each gene is plotted against its mean transcript abundance across all assayed S2 cell samples (cells treated with *stwl* dsRNA and *lacZ* dsRNA as a control). Transcript abundance is represented by counts normalized according to GC-content and library size. The log2(Fold-change) values (LFC) were “shrunk” to minimize the variance associated with low-count genes. Filled points (blue and red) identify genes and repeats which are differentially expressed (adjusted p-value <0.01) in this comparison. Red points represent entries from Repbase, blue points are from the genomic annotation. Y-axis scale is identical to Figure S8, for comparison.

**Figure S12.**
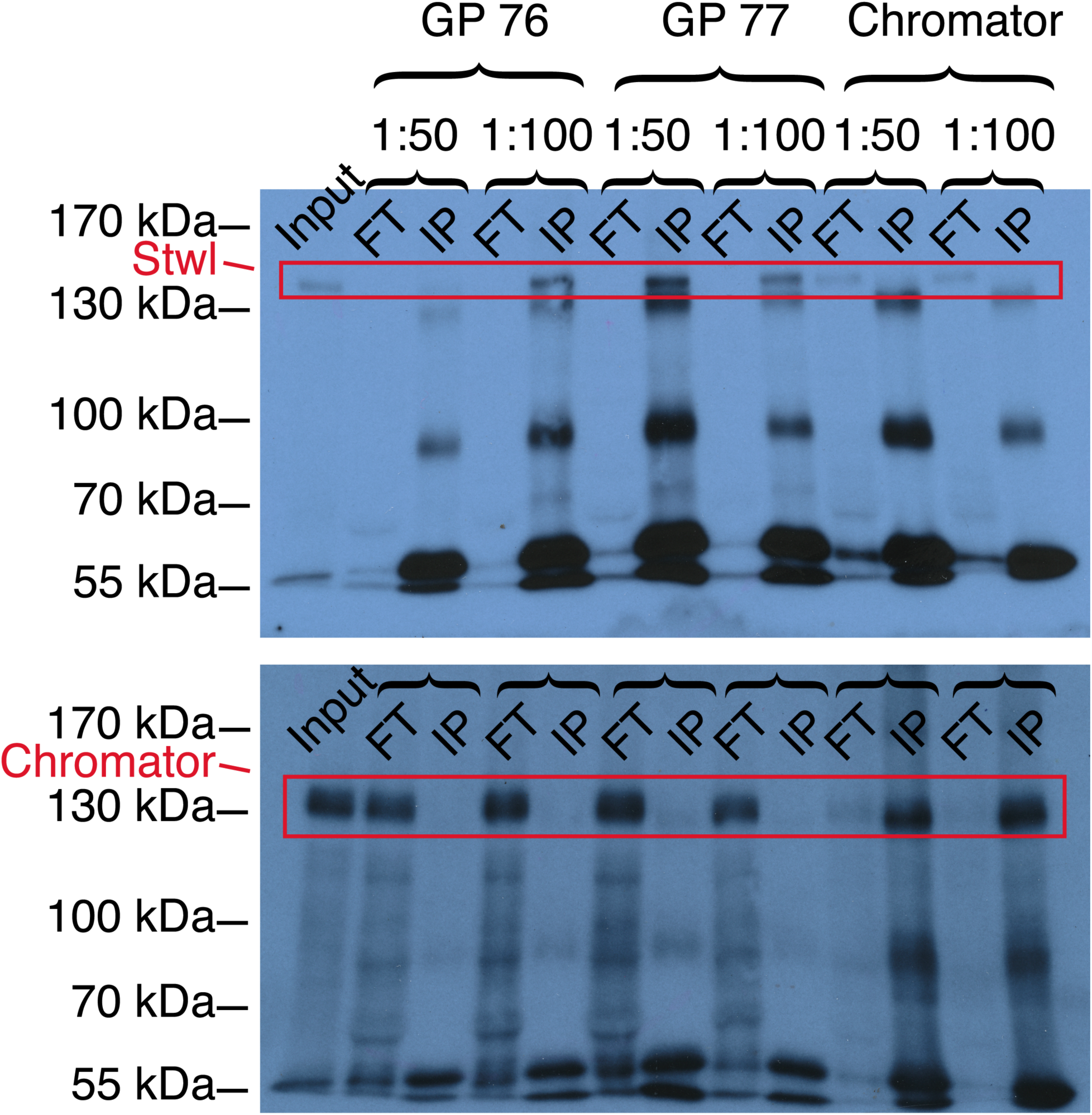
ɑ-Stwl sera immunoprecipitate Stwl from S2 cell lysates. S2 cell nuclei were lysed in RIPA buffer (Input), then incubated with one of two ɑ-Stwl serum or a control antibody (ɑ-Chromator) at 1:50 and 1:100 dilutions. Antibody-Protein complexes were isolated with Protein-A Agarose beads. Western blot of input, flow-through (FT) and IP complexes (IP) probed with ɑ-Stwl GP 76 serum (top panel), then stripped and probed with ɑ-Chromator antibody (bottom panel). Stwl runs at ∼130 kDa (as shown in Figures S1, 3A), as does Chromator. Both ɑ-Stwl sera immunoprecipitate Stwl effectively at a concentration of 1:100 (Stwl protein is eliminated from flowthrough). ɑ-Chromator antibody fails to immunoprecipitate Stwl (Stwl protein remains in flow-through), but successfully immunoprecipitates Chromator.

